# Time-resolved function of cell polarity kinases PRKCZ and PRKCI in CNS myelination

**DOI:** 10.1101/2024.04.22.589759

**Authors:** Maria E. Mercau, Rieke-Marie Hackbarth, Xinran Liu, Haowei Wang, Le Zhang, Malay K. Basu, Carla V. Rothlin, Sourav Ghosh

**Author notes:** These authors contributed equally (Maria E. Mercau, Rieke-Marie Hackbarth). Denotes co-correspondence or.

## Abstract

The atypical Protein Kinase C (aPKC) is a critical component of the PAR polarity complex, as well as of cell polarity in general. There are two paralogs of aPKC in the mammalian genome. Their expression pattern and functional role in oligodendrocyte lineage differentiation and development remains heretofore uncharacterized. Here we show that both paralogs of aPKC – PRKCI, the primary transcript of *Prkci* and PKMz, an alternate transcript of *Prkcz* – are expressed during oligodendrocyte differentiation, yet are functionally non-redundant. Asymmetric cell division and early specification of the oligodendrocyte fate stage required both paralogs. By contrast, the stage of oligodendrocyte maturation involving membrane polarization and myelination of axons continued to require PKMz, but no longer needed PRKCI. Taken together, our results indicate to a two-stage role of aPKCs and cell polarity in oligodendrocyte differentiation and development.

## INTRODUCTION

Cell polarity impacts both oligodendrocyte (OL) differentiation, as well as, functional myelination. Early during OL differentiation, oligodendrocyte progenitor cells (OPC) divide asymmetrically to simultaneously self-renew and give rise to committed oligodendrocyte precursors (COP). This involves cell division of polarized mother cell into non-identical daughters with distinct proliferation *versus* differentiation cell fates ^1,2^. COPs sequentially give rise to newly-formed oligodendrocytes (NFOL) and myelin forming oligodendrocytes (MFOL). Subsequently, MFOLs differentiate into myelinating oligodendrocytes (MOL) ^3^. The myelin-containing membrane of MFOLs and MOLs wraps around the axon sheath of neurons. This myelin membrane is reminiscent of epithelial apical membranes in terms of lipid composition and relative fluidity, by contrast to the plasma membrane surrounding the cell body of the MOL that resembles stereotypical epithelial cell basolateral membranes ^4^. Therefore, the process of myelin membrane formation is also anisotropic. Cell polarity proteins, including atypical Protein Kinase C or aPKC, play a crucial role in the processes of asymmetric cell division and in apical-basolateral membrane polarization ^5,6^. At present, little is known about the role of cell polarity-driven processes during central nervous system (CNS) myelination. Beirowski *et al.* ^7^ showed that lack of histone deacetylase Sirt2 (Sir-two-homolog 2) in Schwann cells impairs PARD3 (Partitioning-defective 3) expression *in vivo*, leading to transient delay in peripheral nervous system (PNS) developmental myelination and remyelination after injury. PARD3 is a component of one of the three major mammalian cell polarity complexes ^8^. The regulation of myelination by PARD3 has been associated with its interaction with p75^NTR^ (neurotrophin receptor p75) at the contact region between Schwann cells and axons. PARD3 absence leads to PNS myelination impairment; however, its downregulation does not affect Schwann cell proliferation or myelin-axon junction formation ^9^. The effect of PARD3 deletion has not been analyzed in the CNS. The Crbs, Pals1/Mpp5 and PatJ complex or Crumbs complex interacts with the PAR complex at the apical membrane ^8^. This complex also contributes to Schwann cell to axon junction formation. Despite no difference in myelin thickness, PALS1 knockout mice presents a reduced conduction of nerve impulses, increased Schmidt-Lanterman incisures frequency and decrease in axon radial sorting. By contrast, PALS1 is dispensable during CNS myelination ^10^, indicating that the myelination process is distinct between PNS and CNS. The third polarity complex – the SCRIB complex, was also implicated in myelination. Jarjour and colleagues have shown that SCRIB is required during OL polarization *in vitro* in response to extracellular matrix ^11^. SCRIB modulated the longitudinal extension of OL, axonal paranodal loop adhesion and remyelination. Daynac *et al.* demonstrated that LGL1 is induced during OL differentiation and conditional knockout of this gene in mice (*Ng2*-cre^ERT2+^ *Lgl* ^f/f^ mice) resulted in impaired OL differentiation ^1^.

The process of cell polarity is considered to be driven by aPKC, a kinase that not only phosphorylates substrates within and thereby regulates the PARD3-PARD6 polarity complex, but also phosphorylates substrates in the Crumbs and SCRIB complexes ^12–14^. In the absence of aPKC, all aspects of cell polarity are thus likely to be severely disrupted, including in oligodendrocytes. For example, *Lgl1*, whose ablation leads to defects in OPC differentiation ^1^, is phosphorylated by and functionally regulated by aPKC ^15^. Despite this *a priori* notion, the functional role of aPKC in oligodendrocyte development and function remains untested. The mammalian genome contains two aPKC genes – *Prkci* and *Prkcz. Prkci* codes for PRKCI (aPKCi) protein, while *Prkcz* encodes two transcripts that are translated into proteins ^14^. The full-length product from transcript 1 of *Prkcz* is PRKCZ (aPKCz). *Prkcz* also encodes an alternative transcript, transcript 2, that gives rise to a shorter, neural isoform termed PKMz ^16,17^. The kinase domains of PRKCZ and PKMz are identical; PKMz lacks only the regulatory domains present in PRKCZ. PRKCI and PRKCZ/PKMz kinase domains are ∼86% identical at the amino acid level ^14^. Therefore, the function of aPKC paralogs may be overlapping and redundant. Nonetheless, previous work from our lab had demonstrated distinct roles for aPKC paralogs PRKCI and PKMz during axon-specification in hippocampal neurons ^18^. PRKCI favored axonogenesis, whereas PKMz suppressed axon formation ^18^. The identity and kinetics of expression of the aPKC paralogs in OL lineage remains undescribed. It is possible that the expression of the aPKC paralogs during oligodendrocyte differentiation is temporally distinct and therefore, relevant to specific cellular events/processes such as OPC to COP differentiation *versus* myelination by MFOL or MOL. Here we examine the time-resolved expression of aPKC paralogs PRKCI, PRKCZ and PKMz in the OL lineage and test their functional requirement in distinct stages of OL differentiation. Conditional knockouts revealed that both paralogs are important for OL development. Specifically, PRKCI and PKMz had functionally non-redundant roles in early OPC differentiation. Additionally, PKMz continued to be required for physiological MOL gene expression and proper myelination, while PRKCI function was dispensable in post-OPC stages.

## RESULTS

### aPKC paralogs *Prkci* and *Prkcz* transcript 2 are expressed in NG2-lineage cells

Interrogation of a previously published study using dissociation and immunopanning to broadly characterize gene expression in OL lineage in postnatal-day (P17) mouse cortices ^19^ revealed that *Prkci* and *Prkcz* are expressed at this stage. RT-PCR analysis demonstrated that *Prkci* (encoding for aPKCι) and *Prkcz* transcript 2 (encoding for PKMζ), but not transcript 1 (encoding for the full-length aPKCζ protein), are expressed in cerebral cortices of C57BL/6 mice at postnatal day (P)30 (**Figure 1A**). *Prkcz* transcript 2 is known to be expressed in the brain, including in the neocortex ^16^. Nonetheless its expression in the white matter was reported to be weak and *Prkcz* transcript 2 was preferentially expressed in neurons ^16^. To test the expression of the aPKC paralogs specifically in OL, we performed RNAscope hybridization in forebrain at P30. OL-lineage cells were identified using a CRE-dependent tdTomato reporter mouse, crossed to an OL-specific CRE-recombinase under the control of the *Cspg* (NG2) promoter ^20^ (*Ng2-cre*^+^ tdTomato ^Tg/+^ mice). RNAscope probes specific for *Prkci* and *Prkcz* transcript 2 were used for hybridization. We observed that both *Prkci* and *Prkcz* transcript 2 were expressed in cerebral cortex OL. Quantification of *Prkci* and *Prkcz* transcript 2-positive puncta in individual OL (tdTomato-positive cells) showed that, at this stage, *Prkcz* transcript 2 was expressed at higher levels than *Prkci* (**Figure 1B**). These results confirm that OL express mRNAs encoding aPKCι and PKMζ.

**Figure 1:**
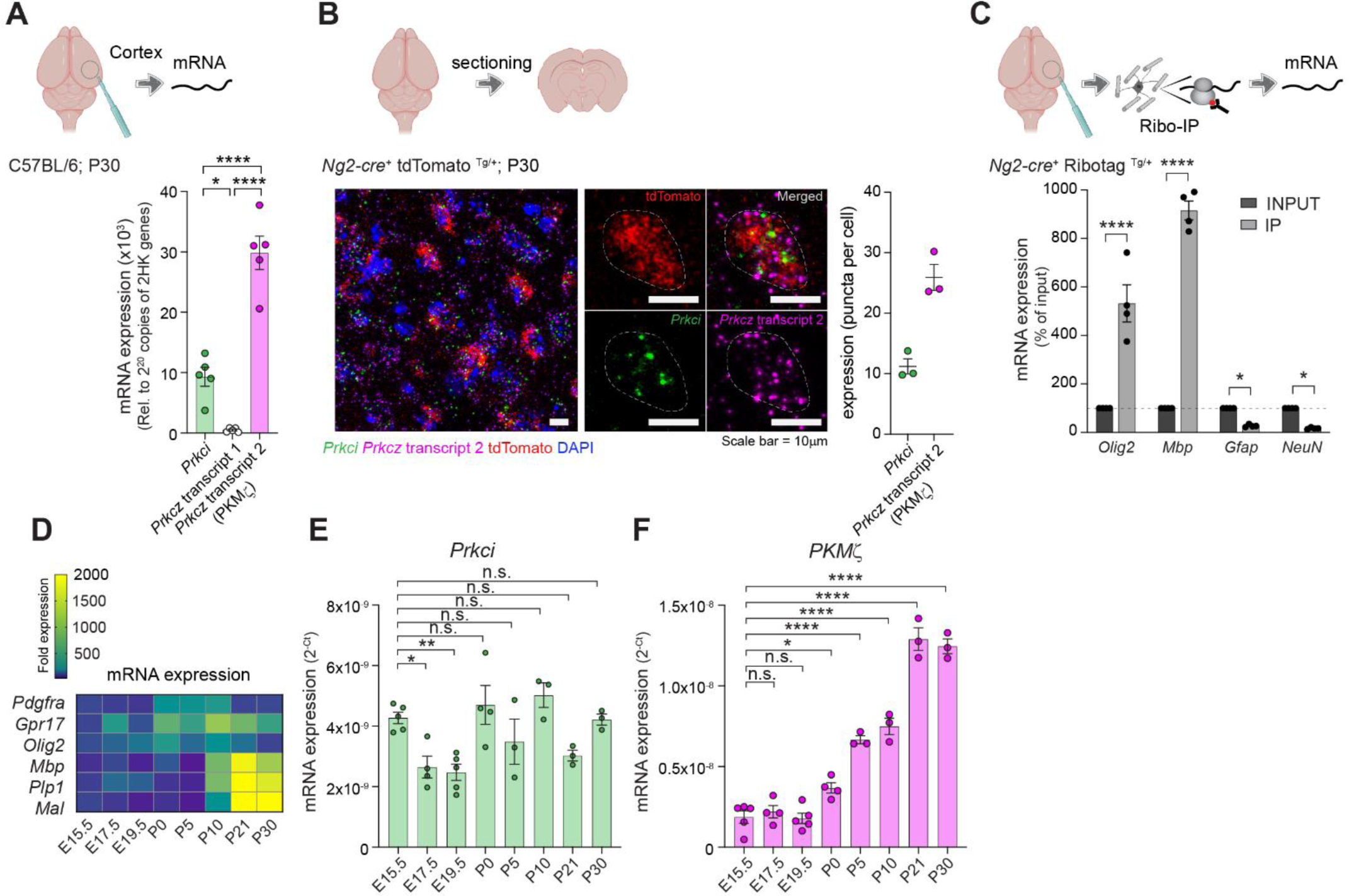
Expression of aPKC paralogs in the brain and in NG2 lineage cells. **A.** mRNA expression of aPKC paralogs in cerebral cortex obtained from C57BL/6 mice at postnatal day (P)30 by qPCR. Data is mean ± SEM, n=5 mice. Non-significant (n.s.), *p<0.05, ****p<0.0001 by ANOVA, followed by Tukey’s test. **B.** aPKC mRNA expression was analyzed in cerebral cortices of P30 *Ng2-cre*^+^ tdTomato^Tg/Tg^ mice by RNAscope. Representative cerebral cortex images are shown, as well as higher magnification images of a single representative oligodendrocyte (tdTomato^+^). Dashed line indicates the area of a single oligodendrocyte. Averages of *Prkci* and *Prkcz* transcript 2 puncta within a single cell were quantified (minimum of 20 oligodendrocytes/ mouse). Data is represented as mean ± SEM, of n=3 *Ng2-cre*^+^ tdTomato^Tg/Tg^ mice. **C.** Cerebral cortices obtained from *Ng2-cre*^+^ Ribotag ^Tg/+^ mice were dissected at the indicated time-points and processed for Ribotag immunoprecipitation. The specificity of ribotag immunoprecipitation was confirmed by analyzing cell type-specific mRNA expression by qPCR in INPUT and IP samples. Data is mean ± SEM, n=4 mice. *p<0.05, ****p<0.0001 vs. INPUT by 2-way ANOVA. **D.** Heatmap of expression of oligodendrocyte stage-specific genes in cerebral cortices dissected at the indicated time-points (starting from embryonic day E15.5 up to P30) from *Ng2-cre*^+^ Ribotag ^Tg/+^ mice. Data is expressed as mean fold expression relative to E15.5. **E.** Expression of *Prkci* was assessed by qPCR in cerebral cortices dissected at the indicated time-points from *Ng2-cre*^+^ Ribotag ^Tg/+^ mice. Data is mean ± SEM, n=3-5 mice/ time point. Non-significant (n.s.), *p<0.05, **p<0.01, *vs.* E15.5 by one way ANOVA, followed by Dunnet’s test. **F.** Expression of *Prkcz* transcript 2 (encoding for PKMζ) was assessed by qPCR in cerebral cortices dissected at the indicated time-points from *Ng2-cre*^+^ Ribotag ^Tg/+^ mice. Data is mean ± SEM, n=3-5 mice/ time point. Non-significant (n.s.), *p<0.05, ****p<0.0001, *vs.* E15.5 by one way ANOVA, followed by Dunnet’s test.

To examine the expression of *Prkci* and *Prkcz* transcript 2 throughout development, specifically in cells from the OL lineage, we utilized mice carrying CRE-recombinase under the control of the *Cspg4* (NG2) promoter as well as a targeted mutation of the ribosomal protein L22 locus that allows CRE-mediated hemagglutinin (HA) epitope tagging of ribosomes (henceforth called Ribotag mice) ^21^. Cortices were collected from *Ng2-cre*^+^ Ribotag ^Tg/+^ mice at different timepoints, and immunoprecipitated ribosome-bound mRNA were subjected to RT-PCR using various gene-specific primers. We first confirmed enrichment of OL-specific genes (*Olig2* and *Mbp*) and depletion of other cell-type-specific genes in the immunoprecipitated fraction as compared to the input (**Figure 1C**). We then analyzed the temporal pattern of expression of OL stage-specific genes to further validate our sample collections (**Figure 1D**). *Mbp*, *Plp1* and *Mal* genes were most highly expressed between P10-P30, while *Pdgfra* was expressed earlier between P0-P10 (**Figure 1D**). Finally, we examined the expression of *Prkci* and *Prkcz* transcript 2. We found that *Prkci* expression was sustained throughout the analyzed period from embryonic day (E)15.5 to P30 (**Figure 1E**), while the mRNA that encodes PKMζ displayed a significant increase during post-natal stages starting at P0, with the highest levels detected at P21 and P30 (**Figure 1F**). In agreement with our RNAscope findings, *Prkcz* transcript 2 was expressed at higher levels compared to *Prkci*, especially at P21-30. These results suggest that the role of aPKCι and PKMζ in cortical OL function may be temporally defined, and therefore, could be related to distinct stages of OL differentiation.

### *Prkci* and *Prkcz* play crucial, non-redundant roles in OL development

To test whether either or both of these isoforms of aPKCs are required for OL development and/or function, we generated mice bearing OL-specific ablation of *Prkci* or *Prkcz*. The targeted genes were ablated by expressing CRE-recombinase under the control of the *Cspg4* (NG2) promoter ^20^ in *Prkci* ^f/f^ ^22^ (**Extended data Figure S1)** or *Prkcz* ^f/f^ ^22,23^ mice. These mice also carried a Ribotag ^21^, which was used to validate OL-specific knockout in *Ng2-cre*^+^ *Prkci* ^f/f^ and *Ng2-cre*^+^ *Prkcz* ^f/f^ mice (**Extended data Figure S2**).

Having confirmed ablation of *Prkci* and *Prkcz* in *Ng2-cre*^+^ *Prkci* ^f/f^ and *Ng2-cre*^+^ *Prkcz* ^f/f^ mice, respectively, we tested the expression of two well-known myelination-associated genes *Mbp* and *Plp1* in cerebral cortices at P30 by RT-PCR (**Figure 2A**). We found a significant reduction in the expression of these genes in samples obtained from *Ng2-cre*^+^ *Prkci* ^f/f^ and *Ng2-cre*^+^ *Prkcz* ^f/f^ mice, relative to control mice (**Figure 2B**). We also measured the expression levels of myelin-associated proteins MBP and MOG by immunoblotting (**Figure 2C**). The levels of myelin proteins were significantly reduced in the cerebral cortices of *Ng2-cre*^+^ *Prkci* ^f/f^ mice (∼38 and 42% reduction of MBP and MOG, respectively) and *Ng2-cre*^+^ *Prkcz* ^f/f^ mice (∼38 and 69% reduction of MBP and MOG, respectively), relative to their corresponding littermate controls (**Figure 2C**). In agreement, luxol fast blue staining intensity was significantly reduced in P45 *Ng2-cre*^+^ *Prkci* ^f/f^ and *Ng2-cre*^+^ *Prkcz* ^f/f^ mice (**Figure 2D**). Taken together, these results demonstrate that both aPKC paralogs *Prkci* and *Prkcz* are required independently for OL development.

**Figure 2:**
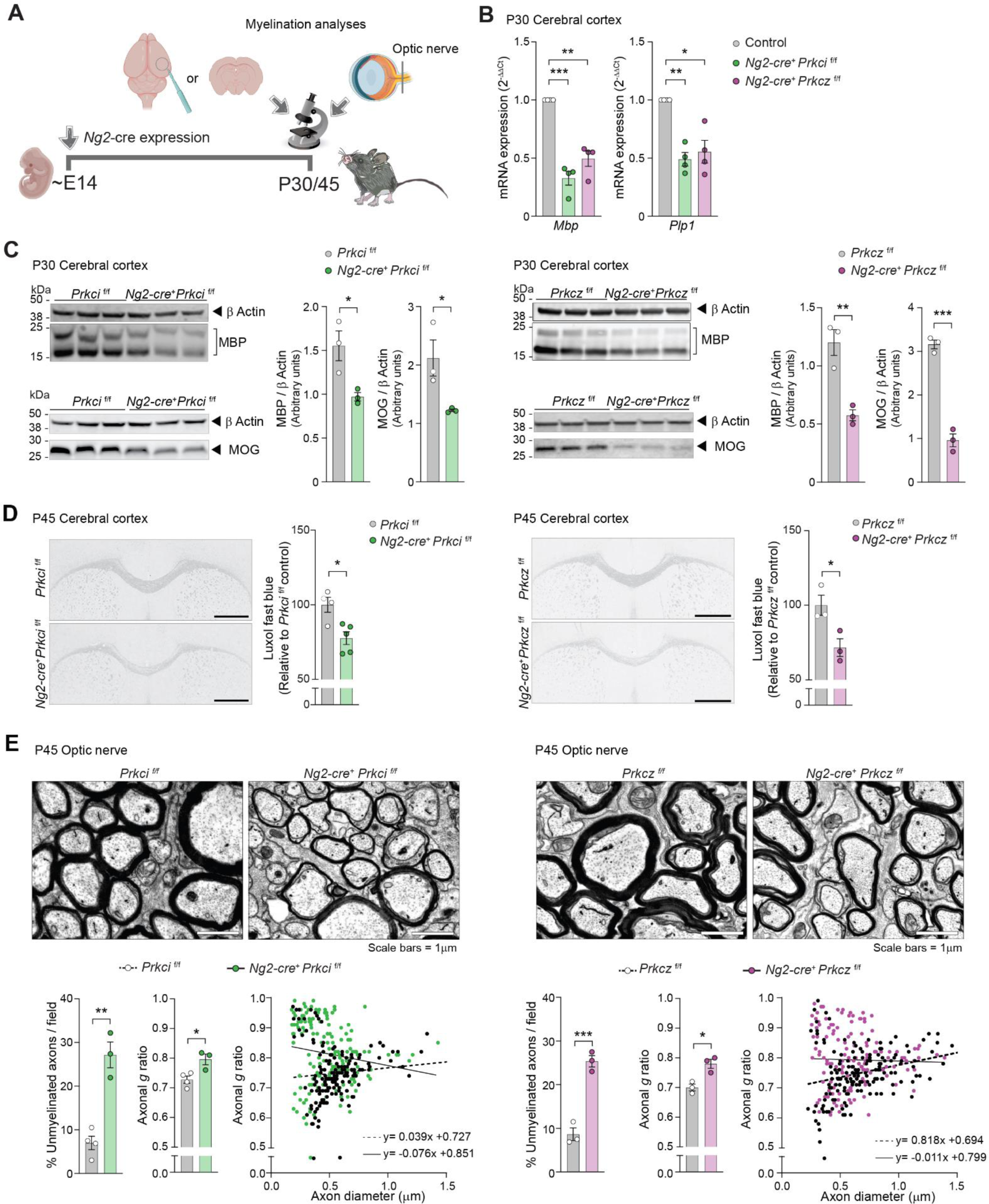
Constitutive genetic ablation of aPKC paralogs *Prkci* or *Prkcz* in NG2 lineage cells results in defective developmental myelination. **A.** Experimental scheme depicting *Ng2-cre* expression at E14 and sample collection and analyses at P30/45. **B.** qPCR for mRNA expression of *Mbp* and *Plp1* in cerebral cortex of P30 mice, normalized to control. Data is mean ± SEM, n=4 mice/ genotype. *p<0.05, **p<0.01, ***p<0.001 by ANOVA, followed by Dunnet’s test. **C.** Cerebral cortices obtained at P30 were dissected and myelin-related proteins were analyzed by western blot. Representative immunoblots for MBP and MOG with corresponding β Actin loading controls, and quantification of *Ng2-cre*^+^ *Prkci* ^f/f^ (left), *Ng2-cre*^+^ *Prkcz* ^f/f^ (right) and littermate controls at P30. Data is mean ± SEM, n=3 mice/ genotype. *p<0.05, **p<0.01, ***p<0.001 by Student’s t test. **D.** Representative images and quantification of luxol fast blue-stained brain sections obtained from P45 *Ng2-cre*^+^ *Prkci* ^f/f^ (left), *Ng2-cre*^+^ *Prkcz* ^f/f^ (right) mice and littermate controls. Data is mean ± SEM, n=3-5 mice/ genotype. *p<0.05 *vs*. control by Student’s t test. **E.** (left) Representative transmission electron microscopy (TEM) images of optic nerves from *Ng2-cre*^+^ *Prkci* ^f/f^ and littermate controls at P45, quantification of % unmyelinated axons in 10 fields/mouse, axonal *g* ratio in 10 fields/mouse and axonal *g* ratio *vs.* axon diameter of optic nerves. Data is mean ± SEM, n=3-4 mice/ genotype. *p<0.05, **p<0.01 by Student’s t test. Linear regression slopes are significantly different (p=0.0186). Scale bars= 1μm. (right) Representative electron microscopy of optic nerves from *Ng2-cre*^+^ *Prkcz* ^f/f^ and littermate controls at P45, quantification of % unmyelinated axons in 10 fields/mouse, axonal *g* ratio in 10 fields/mouse and axonal g ratio *vs*. axon diameter of optic nerves. Axons were determined to be unmyelinated when surrounded by a single layer of membrane/ myelin sheath. Data is mean ± SEM, n=3 mice/ genotype. *p<0.05, ***p<0.001 by Student’s t test. Linear regression slopes are significantly different (p=0.0183). Scale bars= 1μm.

In mice, myelination of the optic nerve occurs along a well-defined time line ^24^. Furthermore, optic nerve myelination is conducive to morphometric analyses of its ultrastructure by transmission electron microscopy (TEM). We ablated *Prkci* or *Prkcz* using *Ng2-cre* and analyzed the effect of genetic ablation of aPKC on optic nerve ultrastructure at P45 (**Figure 2A, E**). Approximately 75% of myelination is completed by P45 and essentially all optic nerve axons are myelinated by P60 ^25,26^, followed by rapid decrease in the rate of myelination ^25,26^ . TEM analysis revealed severe impairment in optic nerve myelination in *Ng2-cre*^+^ *Prkci* ^f/f^ and *Ng2-cre*^+^ *Prkcz* ^f/f^ mice compared to their corresponding littermate controls (**Figure 2E**). Specifically, we found a significant increase in the number of unmyelinated axons per field, a significantly higher average axonal *g* ratio, and decreased myelin thickness in small and medium sized axons (diameter<0.5μm) in optic nerves collected from adult *Ng2-cre*^+^ *Prkci* ^f/f^ and *Ng2-cre*^+^ *Prkcz* ^f/f^ mice compared to their corresponding littermate controls at P45 (**Figure 2E**). Thus, our results on optic nerve myelination phenocopied the effects on cortical myelination and revealed that both aPKC isoforms play a fundamental role in OL function. Importantly, these results also confirmed that these isoforms play non-redundant roles in OL, given that the ablation of one isoform is not compensated by the expression of the other.

### Both aPKC paralogs function in OPC differentiation and early stages of OL development

To understand the effects of aPKC paralogs in OL lineage development, we performed single nucleus (sn)RNA-seq of cerebral cortices from Control and *Prkci* ^f/f^ and *Prkcz* ^f/f^ mice crossed with *Ng2-cre*^+^ or *Plp1*-*cre*^ERT+^ at various time points, using the 10X Genomics platform. Individual transcriptomes from 157,736 nuclei were captured and, based on unbiased clustering, classified into common brain tissue cell types including excitatory neurons (ExN), inhibitory neurons (InN), astrocytes, microglia, endothelial cells, pericytes, vascular and leptomeningeal cells (VLMC), OPC, and mature OL based on well characterized markers (**Figure 3A, B; Extended data Figure S3**). Comparison of Control P30, P60 and P96 time points revealed that the early time point (P30) contained more OPC, COP-NFOL and MFOL while the number of these cell types progressively decreased at P60 and P96 with a concomitant increase in the number of MOLs (**Extended data Figure S4A,B**). Therefore, we tested early OL differentiation *i.e.* peak OPC differentiation into COP-NFOL and MFOLs, at P30. We compared the cellular composition of P30 brain in the presence or absence of aPKC paralogs by genetic ablation using *Ng2-cre*. *Ng2-cre* is expressed early in OL lineage development at E14 ^20^. OL generation peaks between P7-14 in mice ^27^. Therefore, aPKCs are expected to be ablated even before early OPC differentiation into COP-NFOL. In *Ng2-cre*^+^ *Prkci* ^f/f^ mouse brains, the percentage of OL lineage cells and mature OL was reduced by 27% and 36.5%, respectively, by comparison to Control mouse brains (**Figure 3C**). In *Ng2-cre*^+^ *Prkcz* ^f/f^ mice these cells were reduced by 25.8% and 31.9%, respectively by comparison to Control mice (**Figure 3C**). To examine the early differentiation changes in more finely resolved populations of OL lineage cells, we subsetted cells belonging to the OL lineage (OPC and mature OL) and re-clustered them. Proliferating OPC were identified by expression of *Top2a* and *Mki67*. OPCs were defined as *Pdgfr* ^+^ and *Ptprz1 ^+^*, COP/NFOL as *Neu4* ^+^ and *Itpr2* ^+^, MFOL as *Opalin ^+^* and *Ctps* ^+^, and MOL as *Trf* ^+^ and *Apod* ^+^ (**Extended data Figure S4C, D**). The percentage of OPCs, including proliferating OPCs, was increased in both *Ng2-cre*^+^ *Prkci* ^f/f^ (by 35.65%) and *Ng2-cre*^+^ *Prkcz* ^f/f^ mice (by 25.57%) compared to Control mice (**Figure 3D-F**). The percentage of COP-NFOL dropped by 36.70% in *Ng2-cre*^+^ *Prkci* ^f/f^ mice and by 27.34% in *Ng2-cre*^+^ *Prkcz* ^f/f^ mice (**Figure 3D-F**). These results indicate that both *Ng2-cre*^+^ *Prkci* ^f/f^ mice and *Ng2-cre*^+^ *Prkcz* ^f/f^ mice have defects in early OL differentiation. Furthermore, there is no redundancy as ablation of either results in similar differentiation defect.

**Figure 3:**
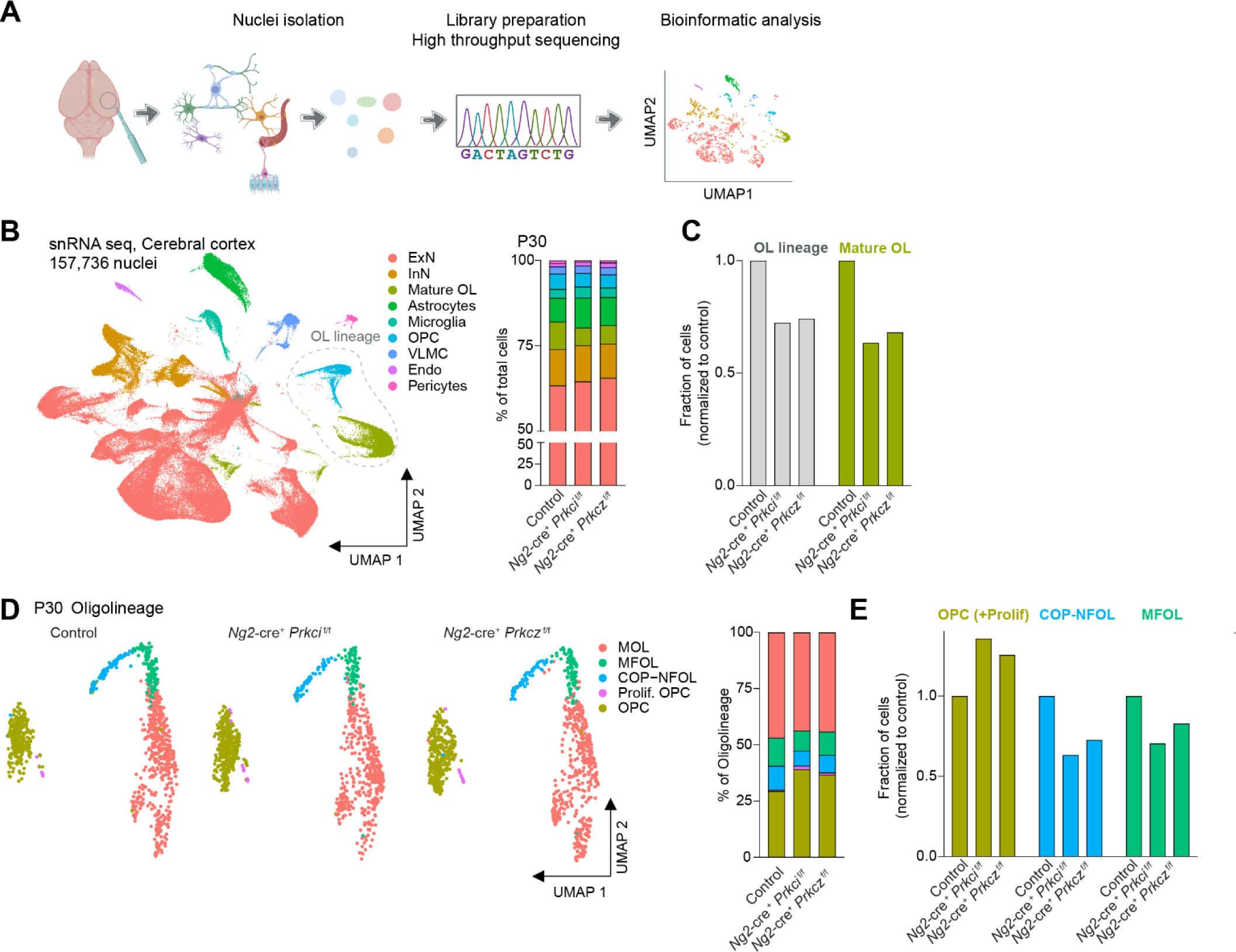
Constitutive genetic ablation of aPKC paralogs *Prkci* or *Prkcz* results in reduced percentage of populations of mature oligodendrocytes. **A.** Schematic of a single-nucleus RNA-Seq (snRNA seq) experiment. Nuclei were isolated from the cortex, RNA sequencing libraries were prepared and sequenced using the 10x Genomics platform, data were then analyzed in R. **B.** Uniform Manifold Approximation and Projection (UMAP) plot of gene expression in 157,736 nuclei from the cerebral cortex, annotated by cell type, determined by snRNA seq. ExN: excitatory neurons, InN: inhibitory neurons, mature OL: mature oligodendrocytes, OPC: oligodendrocytes precursor cells, VLMC: vascular and leptomeningeal cell, Endo: endothelial cells. Dashed line indicates the subset of nuclei used for downstream analysis of oligodendrocyte lineage. Cell type distribution within each sample obtained from Control*, Ng2-cre*^+^ *Prkci* ^f/f^ and *Ng2-cre*^+^ *Prkcz* ^f/f^ at P30 are plotted on the right. **C.** Relative abundance of mature OL and OL lineage in P30 *Ng2-cre*^+^ *Prkci* ^f/f^ and *Ng2-cre*^+^ *Prkcz* ^f/f^ normalized to control. **D.** UMAP of gene expression in single nuclei identified as different stages of oligodendrocyte lineage from Control, *Ng2-cre*^+^ *Prkci* ^f/f^ and *Ng2-cre*^+^ *Prkcz* ^f/f^ mice at P30. MOL: mature oligodendrocytes, MFOL: myelin forming oligodendrocytes, COP: differentiation-committed oligodendrocytes, NFOL: newly formed oligodendrocytes, Prolif. OPC: proliferating oligodendrocyte precursor cells, OPC: oligodendrocyte precursor cells. Percentage of nuclei representing different stages of oligodendrocyte lineage for each genotype are plotted on the right. **E.** Relative abundance of different stages of OL lineage in P30 *Ng2-cre*^+^ *Prkci* ^f/f^ and *Ng2-cre*^+^ *Prkcz* ^f/f^ mice, normalized to control.

We additionally performed bulk RNA sequencing (RNA-seq) of cerebral cortices obtained from Control, *Ng2-cre*^+^ *Prkci* ^f/f^ and *Ng2-cre*^+^ *Prkcz* ^f/f^ mice at P30. Gene set enrichment analysis (GSEA) confirmed reduction in pathways involved in ensheathment of neurons, OL differentiation and development, and glia differentiation as amongst the top downregulated categories in *Ng2-cre*^+^ *Prkci* ^f/f^ and *Ng2-cre*^+^ *Prkcz* ^f/f^ compared to Control mice (**Figure 4A**). Furthermore, expression of selected marker genes corresponding to stages of OL development, such as COP, NFOL, MFOL and MOL^3^ was reduced in *Ng2-cre*^+^ *Prkci* ^f/f^ and *Ng2-cre*^+^ *Prkcz* ^f/f^ mice compared to Control mice (**Figure 4B**). OPC gene expression was not appreciably altered in the absence of *Prkci* or *Prkcz* (**Extended data Figure S5)**. These results validate that the loss of aPKCs independently inhibited proper OPC to COP-NFOL and MFOL differentiation.

**Figure 4:**
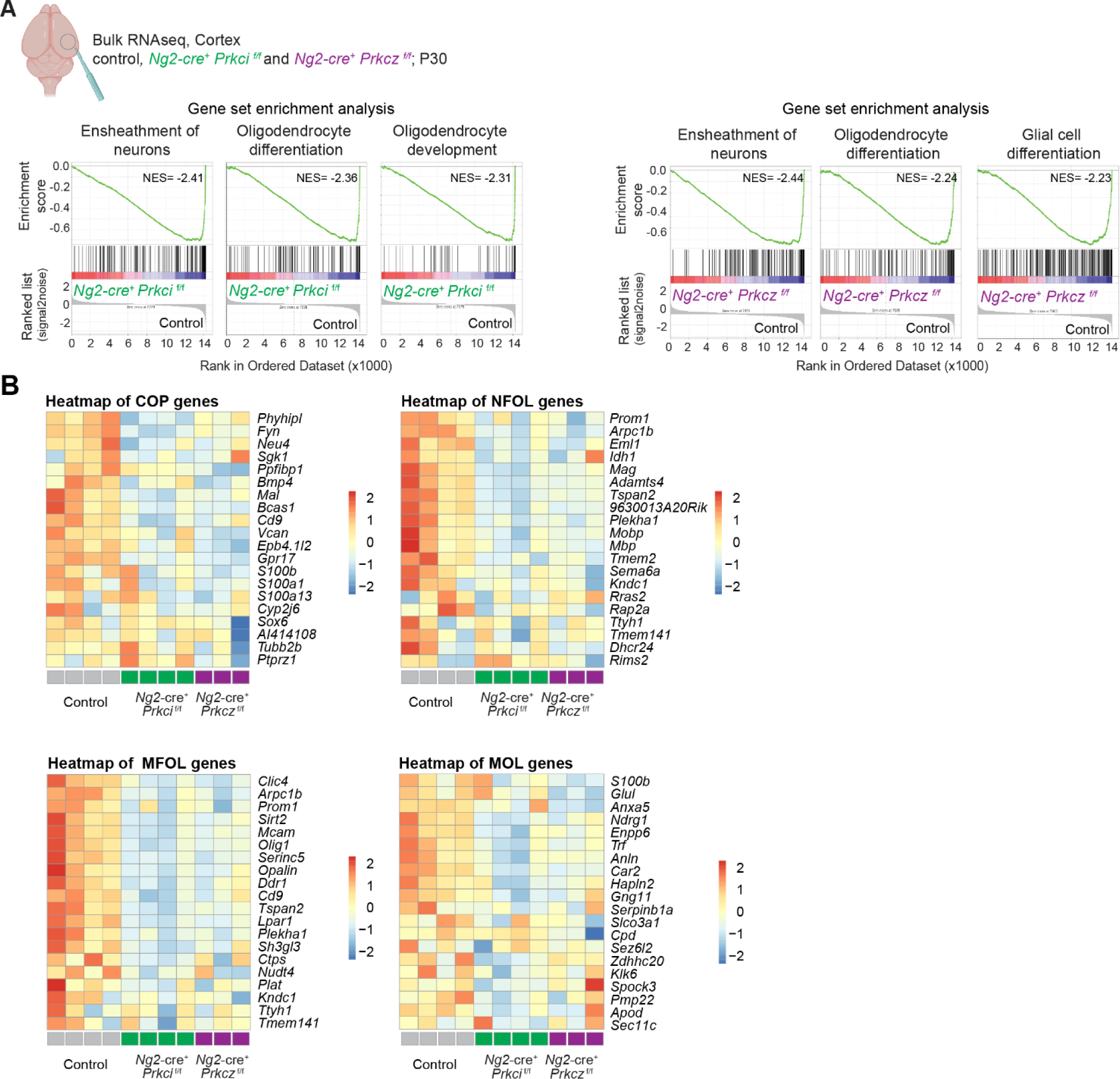
Defective oligodendrocyte precursor to mature oligodendrocyte differentiation in *Ng2*-specific *Prkci* or *Prkcz* mice. **A.** Cerebral cortices obtained at P30 were analyzed by bulk RNA sequencing. Gene-set enrichment analysis (GSEA) of RNA-seq samples from either *Ng2-cre*^+^ *Prkci* ^f/f^ (left) and *Ng2-cre*^+^ *Prkcz* ^f/f^ (right) *vs.* control. Top gene ontology biological process (GOBP) categories are shown. **B.** Heatmap depicting expression of genes expressed by COP, NFOL, MFOL and MOL in indicated samples (based on Marques S. *et al.* 2016). Z-score transformed gene expression values are shown, color-coded according to the color-bar on the right.

### Consistent with its post-natal upregulation, *Prkcz* transcript 2 is indispensable for later stages of OL development

To further test the role of these kinases in later OL developmental stages, we ablated *Prkci* or *Prkcz* in *Plp1*-expressing cells by crossing *Prkci* ^f/f^ and *Prkcz* ^f/f^ mice with mice carrying tamoxifen-inducible CRE recombinase (**Figure 5A)**. Tamoxifen was injected at P35. *Plp1* expression is predominant in post-OPC stages (**Extended data Figure S6**). Furthermore, the bulk of developmental OPC differentiation is completed by P35 ^28^ and myelinating OLs outnumber OPCs at this time point ^27^. We first verified the expression of CRE recombinase 3 weeks after completing tamoxifen dosing using a tdTomato-GFP reporter (mTmG) mouse (**Extended data Figure S7A, B**) and validated the depletion of *Prkci* and *Prkcz* mRNA using *Plp1*-*cre*^ERT+^ *Prkci* ^f/f^ Ribotag ^Tg/+^ and *Plp1*-*cre*^ERT+^ *Prkcz* ^f/f^ Ribotag ^Tg/+^ mice, respectively (**Extended data Figure S7C**). We determined the effects of *Prkci* and *Prkcz* ablation on later stages of OL development by snRNA-seq of cerebral cortices isolated at P96, a time-point with peak amount of MOLs (**Extended data Figure S4**). Interestingly, OL cell composition was significantly different from tamoxifen-treated Control mice only in tamoxifen-treated *Plp1*-*cre*^ERT+^ *Prkcz* ^f/f^, but not in tamoxifen-treated *Plp1*-*cre*^ERT+^ *Prkci* ^f/f^ mice (**Figure 5B**). MOLs were almost entirely absent in tamoxifen-treated *Plp1*-*cre*^ERT+^ *Prkcz* ^f/f^ mice when compared to tamoxifen-treated Control or *Plp1*-*cre*^ERT+^ *Prkci* ^f/f^ mice (**Figure 5B**). *Prkci* function was thus not required after ∼P35-40, post early OL differentiation. This was in sharp contrast to that of *Prkcz*, which was required for the MOL state. In tamoxifen-treated *Plp1*-*cre*^ERT+^ *Prkcz* ^f/f^ mice, a new cellular cluster was present instead of MOLs (**Figure 5B**), likely representing aberrant MOL. In agreement, gene pathways corresponding to ‘ensheathment of neurons’ and ‘galactosylceramide biosynthesis process’ were downregulated in the aberrant MOLs compared to *bona fide* MOLs (**Figure 5C**).

**Figure 5:**
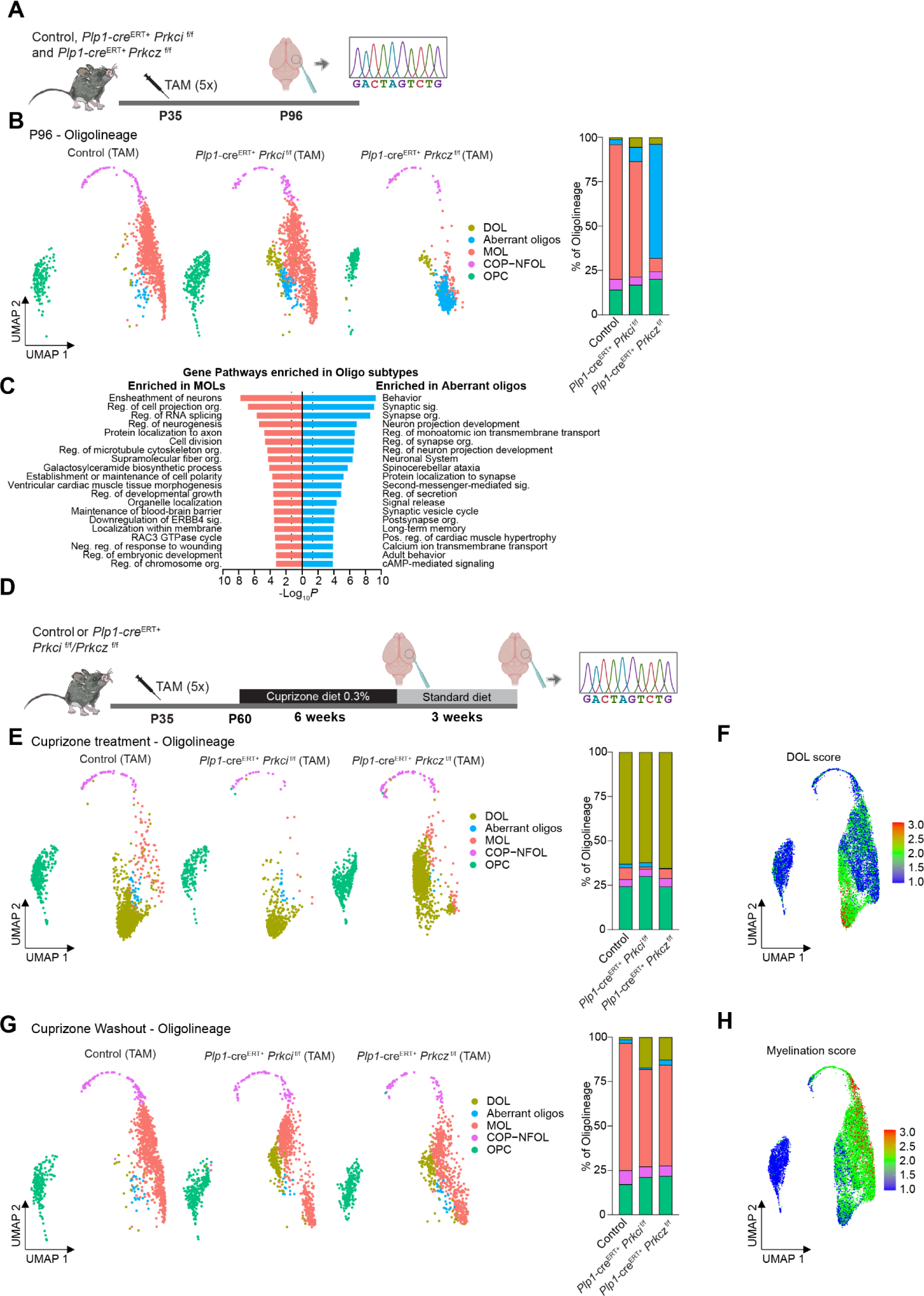
*Prkcz* but not *Prkci* is required for maintenance of myelinating oligodendrocytes in adulthood. **A**. Schematic depicting administration of tamoxifen (5 consecutive days, 1 mg/day, i.p.) to *Plp1-cre*^ERT*+*^ *Prkci* ^f/f^, *Plp1-cre*^ERT*+*^ *Prkcz*^f/f^ and control mice at P35 for 5 days, sample collection and snRNA sequencing performed at P96. **B**. UMAP plot of gene expression in single nuclei classified as different stages of oligodendrocyte lineage from P96 samples. MOL: mature oligodendrocytes, COP: differentiation-committed oligodendrocytes, NFOL: newly formed oligodendrocytes, OPC: oligodendrocyte precursor cells, DOL: demyelination-associated oligodendrocyte. Percentage of nuclei representing different stages of oligodendrocyte lineage for each genotype are plotted on the right. **C**. Enriched gene pathways based on DEGs between MOLs and ‘Aberrant’ oligos. **D**. Experimental schematic depicting *Plp1-cre*^ERT*+*^ *Prkci* ^f/f^, *Plp1-cre*^ERT*+*^ *Prkcz*^f/f^ and control mice injected with tamoxifen at P35 and subsequently fed with 0.3% cuprizone for 6 weeks. In another group, 6 weeks of cuprizone treatment were followed by 3 weeks of standard diet (Cuprizone Washout group). **E**. UMAP plot of gene expression in single nuclei classified as different stages of oligodendrocyte lineage from cuprizone-treated samples separated by genotype. Percentage of nuclei representing different stages of oligodendrocyte lineage for each genotype are plotted on the right. **F**. Feature plot of DOL genes score in oligodendrocyte lineage populations. Genes used to generate the score were previously defined by Hou *et al.* 2023. **G**. UMAP plot of gene expression in single nuclei classified as different stages of oligodendrocyte lineage from Cuprizone Washout samples. Percentage of nuclei representing different stages of oligodendrocyte lineage are plotted on the right. **H**. Feature plot of Myelination genes score in oligodendrocyte lineage populations. Gene signatures related to myelination were derived from the ’Myelination’ pathway as identified through Metascape pathway analysis.

It has been described that cuprizone treatment in mice induces MOLs to undergo transition to a demyelination-associated oligodendrocyte (DOL) state ^29^. To determine if indeed the ‘aberrant MOL’ population is MOL-like and can undergo this transition into DOLs, we administered cuprizone to tamoxifen-treated *Plp1*-*cre*^ERT+^ *Prkci* ^f/f^ or *Plp1*-*cre*^ERT+^ *Prkcz* ^f/f^ mice (**Figure 5D**). 6 weeks of feeding mice with pellets containing 0.3% cuprizone resulted in the *bona fide* MOL in tamoxifen-treated Control and *Plp1*-*cre*^ERT+^ *Prkci* ^f/f^ mice, as well as the ‘aberrant MOL’ in tamoxifen-treated *Plp1*-*cre*^ERT+^ *Prkcz* ^f/f^ mice, to be replaced by a cluster with the molecular signature of DOLs (**Figure 5E, F**). Therefore, the differentiated, aberrant, OL cells observed in tamoxifen-treated *Plp1*-*cre*^ERT+^ *Prkcz* ^f/f^ mice are capable of transitioning into DOLs in response to cuprizone, suggesting that these cells are indeed MOL-like. To verify that the ‘aberrant MOLs’ result from the absence of *Prkcz*, we washed out cuprizone for 3 weeks to observe whether *bona fide* MOLs can be regenerated in *Plp1*-*cre*^ERT+^ *Prkcz* ^f/f^ mice. After cuprizone washout, OPCs repopulate MOLs for remyelination. Importantly, no additional tamoxifen was injected during this period of remyelination, and *Plp1*-*cre*^ERT+^ is not expressed in OPCs. Therefore, tamoxifen treatment between P35 to P40 did not target the OPCs that repopulate MOL post cuprizone-induced demyelination. Cuprizone washout and remyelination in *Plp1*-*cre*^ERT+^ *Prkcz* ^f/f^ mice led to a recovery of *bona fide* MOL cluster, with a concordant myelination molecular signature, in these mice (**Figure 5G, H**). Taken together, our results indicate that it is the absence of *Prkcz* that results in aberrant MOLs, suggesting that PKMz is required for formation and/or maintenance of a *bona fide* MOL state.

### Temporally distinct requirement of aPKC paralogs for optic nerve myelination

To test if the paralogs have a similar time-resolved function during optic nerve myelination, we again took advantage of tamoxifen-inducible activation of CRE recombinase in *Plp1*-*cre*^ERT+^ *Prkci* ^f/f^ and *Plp1*-*cre*^ERT+^ *Prkcz* ^f/f^ mice (**Figure 6A**). *Plp1*-*cre*^ERT+^ mediated ablation of *Prkci* (*Plp1*-*cre*^ERT+^ *Prkci* ^f/f^ mice) was induced starting at P15 by 5 successive days of tamoxifen injections. Optic nerve myelination was analyzed by TEM at P60. By contrast to the effects of *Ng2-cre* induced ablation of either isoform, the deletion of *Prkci* after P15 revealed no significant changes in the number of unmyelinated axons and in the axonal *g* ratio (**Figure 6B**). Thus, *Prkci* function is required exclusively in an early time window, even for optic nerve myelination. By contrast, when *Prkcz* was ablated by the same approach of 5 successive days of tamoxifen injections starting at P15, we observed significantly increased number of unmyelinated axons and increased axonal *g* ratio, specifically in axons with diameter <0.5mm (**Figure 6B**), indicating an impairment in optic nerve myelination in these mice. To further investigate the timeframe for the requirement of *Prkcz* in myelination, we injected Control and *Plp1*-*cre*^ERT+^ *Prkcz* ^f/f^ mice with tamoxifen starting at P25 or P35 (**Extended data Figure S8A, C**) and analyzed the ultrastructure of their optic nerves at P60 (**Extended data Figure S8B, D**). We found that *Plp1*-*cre*^ERT+^ *Prkcz* ^f/f^ mice injected with tamoxifen starting at P25 developed myelination deficits, as evidenced by significantly increased number of unmyelinated axons and axonal *g* ratio, as compared to their littermate controls (**Extended data Figure S8B**). However, *Plp1*-*cre*^ERT+^ *Prkcz* ^f/f^ mice injected with tamoxifen starting at P35 no longer showed significant changes in these parameters (**Extended data Figure S8D**), indicating that *Prkcz* expression becomes dispensable after P35, likely because myelination of the optic nerve is complete/almost complete by this time. Taken together, these results demonstrate that, analogous to that observed in cortical myelination, *Prkci* functions early during optic myelination nerve while *Prkcz* is required till a later time point. These results broadly agree with our findings on cortical myelination (**Figure 2-5**).

**Figure 6:**
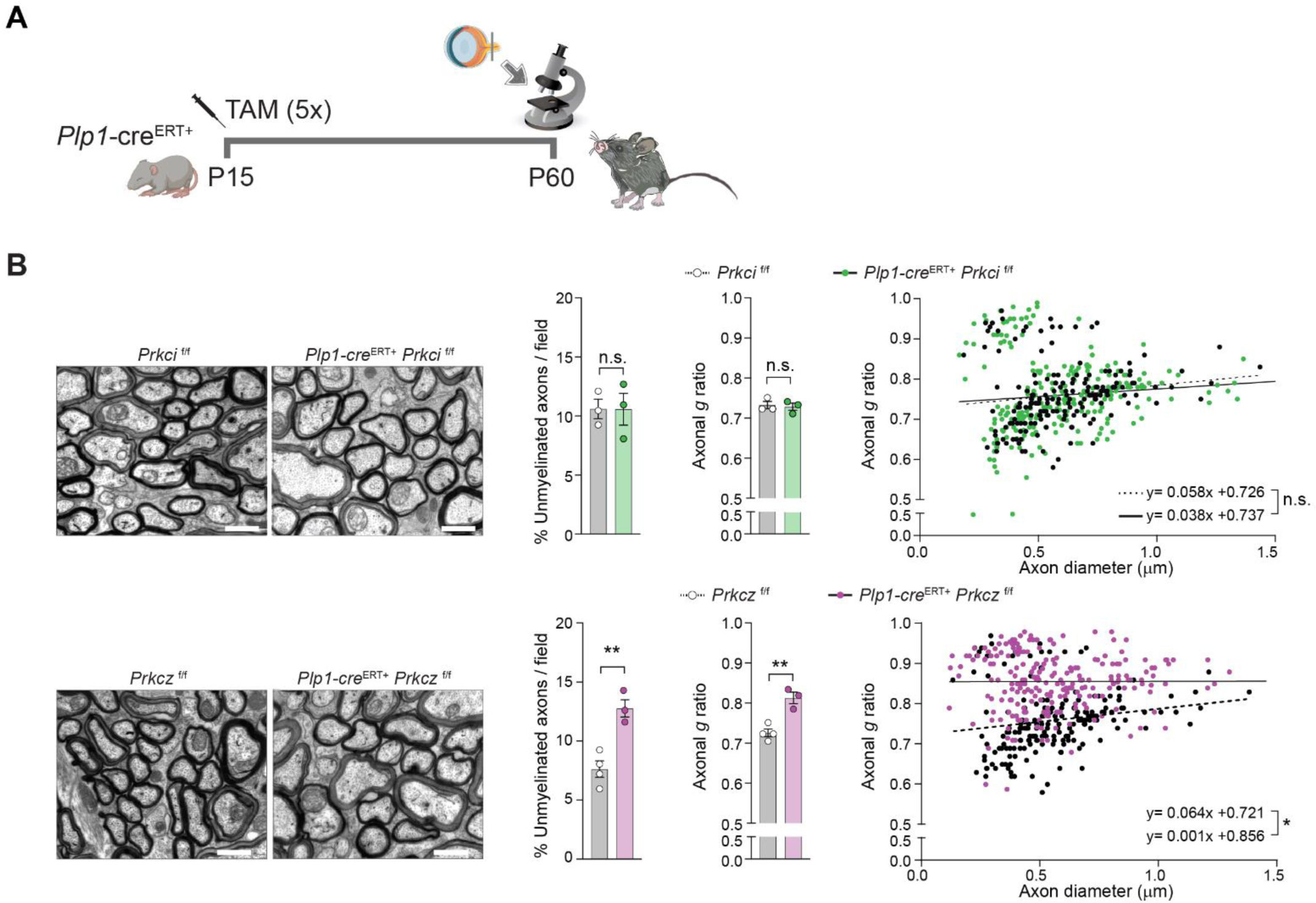
*Prkcz* but not *Prkci* is required at later stages of myelination of the optic nerve. **A.** Schematic depicting administration of tamoxifen (5 consecutive days, 1 mg/day, i.p.) to *Plp1-cre*^ERT*+*^ *Prkci* ^f/f^, *Plp1-cre*^ERT*+*^ *Prkcz*^f/f^ and control mice at P15 for 5 days, sample collection was performed at P60. **B.** (top) Tamoxifen was administered to *Prkci* ^f/f^ and *Plp-cre*^ERT+^ *Prkci* ^f/f^ mice (5 consecutive days, 1 mg/day, i.p.) starting at P15. Mice were sacrificed at P60, and optic nerves were analyzed by TEM. Representative TEM images are shown next to quantification of % unmyelinated axons per field in 10 fields/mouse, axonal *g* ratio in 10 fields/mouse and axonal *g* ratio *vs*. axon diameter. Data is represented as mean ± SEM, n=3 mice/ genotype. Non-significant (n.s.) by Student’s t test. Linear regression slopes are not significantly different (p=0.6124). Scale bars = 1μm. (bottom) Tamoxifen was administered to *Prkcz* ^f/f^ or *Plp-cre*^ERT+^ *Prkcz* ^f/f^ mice (5 consecutive days, 1 mg/day, i.p.) starting at P15. Mice were sacrificed at P60, and optic nerves were analyzed by TEM. Representative TEM images are shown next to quantification of % unmyelinated axons per field in 10 fields/mouse, axonal *g* ratio in 10 fields/mouse and axonal *g* ratio *vs*. axon diameter. Axons were determined to be unmyelinated when surrounded by a single layer of membrane/ myelin sheath. Data is mean ± SEM, n=3-4 mice/ genotype. **p<0.01 by Student’s t test. Linear regression slopes are significantly different (p= 0.0459). Scale bars = 1μm.

## DISCUSSION

We demonstrate that two paralogs of the cell polarity kinase, aPKC, are expressed during OL development. *Prkci* is expressed at steady levels during E15.5 to P30. The transcript that is expressed from *Prkcz* is transcript 2 – its mRNA encodes PKMz. PKMz lacks the regulatory domain of full-length aPKC protein, but can still bind the polarity scaffold PARD3 ^18^. It was previously shown to be important for neuronal polarization ^18^. Our results also implicate *Prkci* and *Prkcz* in the development and function of OL. The loss of either paralog exhibits a myelination defect, indicating that these paralogs are not merely redundant.

Both aPKC paralogs are required at early stages of OL development *i.e.* OPC differentiation into COP-NFOL. The specific mechanistic role of aPKC paralogs in early OL development *i.e.* OPC differentiation is not known. If both paralogs simultaneously function at the stage of OPC differentiation, yet are non-redundant, one possible model may be of competition between the paralogs to establish cell polarity and asymmetric cell division. We previously described this model for axon *versus* dendrite formation ^18^. The loss of either paralog would abolish competition and therefore function. Consistent with a role of these kinases in asymmetric cell division, we noticed a skewing of post-neuronal differentiation of glia away from OL lineage towards astrocytes when *Prkci* and *Prkcz* were ablated using *Ng2*-cre (**Figure 3B**). *Ng2* is expressed in glial common progenitors that give rise to astrocytes and OL ^20^. An alternative model may be that the kinases phosphorylate a unique set of substrates, despite sharing ∼86% amino acid identity in their kinase domain. This may occur, for example, as PKMz lacks the N-terminal region present in PRKCI and consequently the PARD6 binding PB1 domain. Thus, any substrate recruitment via PB1 domain will not happen for PKMz. Furthermore, since PKMz lacks the regulatory domain, its substrate binding in the kinase domain may be of a higher affinity/unconstrained by the regulatory domain and thus differ from PRKCI. Nevertheless, downstream signaling from both sets of substrates has to be critical for OPC differentiation.

Differentiation of OPC involves asymmetric division of OPC to give rise to COP. COP differentiates into NFOL before transitioning into myelinating OL. In the absence of *Prkci* and *Prkcz* transcript 2, there is an increased abundance of OPC. Whether this is due to increased symmetric as opposed to asymmetric divisions within OPC as a direct result of loss of polarity signaling kinase or due to compensation as a result of reduction in MOL remains unclear at this time. An increase in OPC numbers even after ablation of *Prkcz* in later stages (post P35) by using tamoxifen-induced *Plp1*-*cre*^ERT+^ suggests that the increase in OPC, at least in part, may be due to compensatory mechanisms as MOL numbers are reduced. Increase in OPC number is also observed after cuprizone-induced ablation of MOL, further supporting this concept. Notwithstanding, it is likely that the aPKC paralogs are critical for asymmetric cell division.

*Prkcz* transcript 2 is upregulated during postnatal development in mice. NFOL differentiates into MFOL and subsequently MOL. Maturation of OL and myelination also involves specialization of a liquid-ordered myelin membrane that contains approximately 70% of its dry weight in lipids such as cholesterol, galactolipids, galactosylceramide and sulfatide and specialized proteins such as PLP, MBP and MOG. The myelin membrane bears resemblance to apical membrane of epithelial cells and is distinct from liquid-disordered plasma membrane that resembles basolateral membrane of epithelial cells ^4^. While *Prkci* is dispensable for generation of MOLs, *Prkcz* continues to be critical. PKMz loss compromises MOL identity. The downregulation of gene pathways for ‘ensheathment of neurons’ and ‘galactosylceramide biosynthesis process’ in aberrant MOLs suggest that PKMz is important for specialization of the myelin membrane. Consistent with a role in physiological myelination, *Prkcz* locus has been shown to be epigenetically silenced in brains from multiple sclerosis patients ^30^.

What is the mechanism of PKMz function in late OL development? The upregulation in *Prkcz* transcript 2 during postnatal development may offset competition between PRKCI and PKMz, if this indeed is the mechanism of aPKC function in early development. Thus, at later stages of OL development, PKMz function is distinct from its early molecular function of antagonizing PRKCI. Notably, only PKMz continues to function in late OL development in both cortical myelination and in optic nerve myelination, although there are critical differences. We observed aberrant MOLs when *Prkcz* was ablated late in the cortex (P35-40), but no longer observed a dependance on *Prkcz* in optic nerve myelination. It is possible that PKMz has distinct functions in the cortex *versus* the optic nerve, but it is likely that PKMz is required for MOL differentiation and not for MOL maintenance. OPC differentiation continues in the cortex throughout life ^27,31–33^ and therefore the effect of *Prkcz* ablation would be evident in this tissue at P35-40. By contrast, myelination is largely completed in the optic nerve by this time point ^24^. Thus, any effect of *Prkcz* ablation on OPC differentiation is likely to be marginal by this time point. An effect on maintenance, contrary to that in differentiation, can be expected to be observed in the optic nerve at P35-40. In conclusion, two distinct paralogs of mammalian aPKCs are expressed and required for CNS myelination. Furthermore, while both paralogs are required for OPC differentiation, PRKCI is expendable for later stages of OL development, while PKMz continues to be critical for MOL differentiation.

## Acknowledgements

This work was supported by Yale Funds (C.V.R. and S.G.). RM. H. was funded by Boehringer Ingelheim Fonds. We thank the Yale Center for Research Computing for guidance and use of the research computing infrastructure. We thank the Yale Center for Genome analysis and acknowledge that research reported in this publication was supported by the National Institute of General Medical Sciences of the National Institutes of Health under Award Number 1S10OD030363-01A1. Figures were created with BioRender.com.

## Materials and Methods

### Animals

Mice were bred and housed in a pathogen-free facility under standard conditions (12:12 light:dark cycle, with unrestricted access to food and water) and handled in accordance with the guidelines set by the Institutional Animal Care and Use Committee at Yale University and Yale Animal Resources Center. All the mouse strains used were on a C57BL/6 background. Both male and female mice were used in experiments. All efforts were made to minimize the pain and the number of animals used.

### mRNA isolation and quantitative RT-PCR

For total mRNA isolation, mice were euthanized at the indicated postnatal timepoints by CO_2_ inhalation and brains were immediately dissected on ice. Equal amounts of tissue were obtained from cerebral cortices using a 4mm punch biopsy tool, flash frozen in liquid nitrogen and stored at -80°C until further processing. Total mRNA was extracted using RNAeasy Mini Kit, as per manufacturer’s instructions (QIAGEN). Reverse transcription was carried out using iScript cDNA synthesis kit (BioRad), and quantitative (q)PCR reactions were carried out using CFX96 Touch Real-Time PCR Detection System (BioRad). A list of oligonucleotides used for this study is included in Table S1.

### Ribo-qPCR

For Ribotag experiments, lineage-specific HA-tagging of ribosomal subunits was achieved by crossing RiboTag mice ^21^ to either *Ng2-cre*^+^ or *Plp1-cre*^ERT+^ cre-recombinase drivers. Mice were sacrificed at the corresponding postnatal time point by CO_2_ inhalation. Brains were immediately dissected, and 4mm biopsies from cerebral cortices were flash frozen in liquid nitrogen and stored at -80°C until further processing. Ribotag immunoprecipitation and RNA isolation were performed following previously published protocols^34^. Briefly, 200μl of Dynabeads were washed and incubated in citrate-phosphate buffer with 10μl of anti-HA antibody for 45 min at 4°C on a rotating platform. Cerebral cortices of 2 mice per time point were pooled and homogenized in homogenization buffer containing cycloheximide and RNAse inhibitors at a concentration of 10% m/v. After centrifugation at 10000xg for 10 min, 10% of the sample was dissolved in RLT buffer and frozen for further processing of INPUT. The remaining sample was incubated with the antibody-coupled beads on a rotating platform at 4°C for 6 hours. After washing, the beads containing the immunoprecipitated (IP) samples were washed and incubated with RLT buffer for 10 minutes on a thermomixer (750rpm) at room temperature. Supernatants were collected and RNA was obtained from the IP and INPUT samples using QIAGEN RNeasy kits (QIAGEN). Reverse transcription was carried out using iScript cDNA synthesis kit (BioRad), and PCR reactions were carried out using CFX96 Touch Real-Time PCR Detection System (BioRad).

### Bulk RNA sequencing

Library preparation and sequencing of RNA from cerebral cortex samples (3-4 biological replicates for each genotype) was performed at the Yale Center for Genome Analysis using 100-base pair, paired-end reading on a NovaSeq instrument (Illumina) at a sequencing depth of 25 million reads per sample. Raw reads were input into FastQC ^35^ to perform initial quality control checks and TrimGalore was used to trim adapter sequences. We employed STAR ^36^ for alignment of the trimmed reads to the annotated mouse genome GRCm38. Transcript abundance was quantified using RSEM ^37^. GSEA^38^ was employed to evaluate whether defined sets of genes associated with specific pathways show statistically significant, concordant differences between two biological states. Pathways were considered significantly enriched based on high normalized enrichment score and an FDR < 0.25. Heatmaps depicting z scores for top 20 genes (Oligodendrocyte precursor cells, differentiation-committed oligodendrocytes, myelin forming oligodendrocytes, mature oligodendrocytes, from Marques *et al.,* 2016) were generated using the pheatmap package in R.

### Western blot

Mice were sacrificed at the corresponding postnatal time point by CO_2_ inhalation. Brains were dissected and 4mm punch biopsies of cerebral cortices were flash frozen in liquid nitrogen and stored at -80°C until further processing. Samples were homogenized in RIPA buffer containing cOmplete™ Protease Inhibitor Cocktail (Roche). Protein concentration was determined by spectrophotometry using BCA assay (Thermofisher). For immunoblots, equal amounts (30μg) of total protein in Laemmli Buffer were subjected to electrophoresis on precast polyacrylamide gels and transferred to PVDF membranes (BioRad). Membranes were blocked and probed overnight with primary antibodies. Secondary antibodies conjugated to near infrared fluorophores (Li-COR Biosciences Cat #926-32213 and Cat #926-68072) were detected using Odyssey Imaging System and quantified with Image Studio Lite Software.

### Transmission Electron microscopy

Adult mice were intracardially perfused with PBS followed by 4% paraformaldehyde (PFA) in 1x PBS, optic nerves were then carefully dissected, and further fixed in 2.5% glutaraldehyde and 2% PFA in 0.1 M sodium cacodylate buffer (pH 7.4) for one hour at room temperature. They were post-fixed in 1% OsO4, 0.8% potassium ferricyanide in the same buffer for one hour at room temperature, then *en bloc* stained with 2% aqueous uranyl acetate for 30 min. Optic nerves were then dehydrated in a graded series of ethanol to 100%, substituted with propylene oxide, embedded in Embed 812 resin, and polymerized in an oven at 60 °C overnight. Optic nerves were cut 1-2 mm proximal to the optic chiasm into thin sections of 60 nm by a Leica ultramicrotome (UC7), placed on standard EM grids and stained with 2% uranyl acetate and lead citrate. Samples were examined with a FEI Tecnai transmission electron microscope at 80 kV accelerating voltage, digital images were recorded with an Olympus Morada CCD camera and iTEM imaging software at the Yale Center for Cellular and Molecular Imaging (CCMI) Electron Microscopy Facility. Images were analyzed manually using ImageJ software. For each axon, the thickness of the surrounding myelin (Δi) was calculated as the average of manual measurements made in two locations, and the *g*-ratio was estimated as g = r/(r + Δi). Axons were determined to be unmyelinated when surrounded by a single layer of membrane/ myelin sheath.

### Histology/anatomical studies

#### RNAscope in situ hybridization

Tools, slides, and equipment were cleaned with 70% ethanol and ribonuclease inhibitors (Fisher Scientific). Mice were euthanized by cervical dislocation, and brains were rapidly dissected, and flash frozen with 2-methylbutane chilled on dry ice for 20 seconds. Brains were subsequently stored at −80°C until sectioning. Brains were sliced using a cryostat (Leica), and sections (16 μm) were mounted on charged microscope slides. RNAscope (Advanced Cell Diagnostics) *in situ* hybridization procedures were conducted according to the manufacturer’s instructions. Briefly, slides were fixed with 10% neutral buffered formalin for 20 min at 4°C, washed twice for 1 min with 1x PBS, before dehydration with increasing percentages of ethanol. Slides were then stored in 100% ethanol at −20°C overnight. The next day, slides were dried at room temperature (RT) for 10 min and incubated with Protease Pretreat-4 solution for 20 min at RT. Slides were washed twice with ddH2O before being incubated with the appropriate probes for 2 hours in a humid chamber at 40°C. Probes used were purchased from Advanced Cell Diagnostics as follows: tdTomato (317041-C2), Mm-*Prkci* (403191-C1), and Mm-*Prkcz* (403201-C3). Images were acquired using a Zeiss LSM 800 series confocal microscope at the Yale CCMI facility, with an oil-immersion 63X objective.

#### Luxol fast blue staining

Luxol fast blue (LFB) staining was performed by the Yale Pathology Tissue Services facility at Yale School of Medicine following standard procedures. Briefly, adult mice were intracardially perfused with 1x PBS followed by 4% PFA, brains were then carefully dissected, and prepared for paraffin embedding. 6 μm sections were cut, deparaffinized, hydrated, and subsequently stained in 0.1% LFB solution overnight at 50°C. After differentiation with 0.05% lithium carbonate, sections were dehydrated in increasing ethanol concentrations and mounted with resinous mounting medium (Permount^®^).

#### Immunofluorescence

Animals were sacrificed at the indicated time points by CO_2_ inhalation and brains were immediately dissected and fixed in 4% PFA solution overnight. After 30% sucrose dehydration, brains were embedded in OCT and cut in a cryostat (Leica Biosystems) to a thickness of 30μm. Antigen retrieval was performed using citrate phosphate buffer (pH 6) at 80°C for 30 min. After blocking in 5% normal goat serum in 1% PBS-T, samples were stained with primary antibodies anti-GFP overnight at 4°C following a free-floating protocol. After washing, samples were incubated with Alexa Fluor^®^-conjugated secondary antibodies for 1 hour at room temperature and mounted on slides using DAPI-containing mounting media (Dako). Images were acquired using a Zeiss LSM 800 series confocal microscope at the Yale CCMI facility, with an oil-immersion 10X objective. Images were analyzed using ImageJ software.

### Tamoxifen-induced ablation

*Plp1*-*cre*^ERT^ mice (Jackson Laboratories stock no. 005975) were crossed to either *Prkci* ^f/f^ or *Prkcz* ^f/f^ mouse lines. A 10-mg/ml tamoxifen stock suspension was prepared for the induction experiment by mixing 10 mg tamoxifen with 100μl of ethanol and 900 μl of sunflower oil, followed by 30 min sonication. Mice were injected at the indicated time points for 5 consecutive days intraperitoneally with 100 μl of suspension (1mg tamoxifen/day), as previously described ^39^.

### Cuprizone treatment

*Plp1-cre*^ERT+^ *Prkci* ^f/f^, *Plp1-cre*^ERT+^ *Prkcz* ^f/f^ mice and littermate controls were injected with 4-OH Tamoxifen (Sigma Aldrich, MO) for 5 days starting at postnatal day 35. Two weeks after tamoxifen delivery, cuprizone-mediated demyelination was induced by feeding mice pellets containing 0.3% cuprizone (Envigo, IN) for 6 weeks. Food was refreshed and mice were monitored daily. For remyelination studies (cuprizone washout group), food was switched to standard diet for 3 weeks prior to sample collection.

### Mouse brain nuclei isolation and single nucleus RNA-sequencing

Nuclei were isolated from flash-frozen mouse cerebral cortices collected at indicated postnatal days from indicated genotypes (6mm punch biopsies from n=3-5 mice of each genotype were pooled per sample). 15 ml of ice-cold nuclei homogenization buffer [2 M sucrose, 10 mM Hepes (pH 7.9), 25 mM KCl, 1 mM EDTA (pH 8.0), 10% glycerol, and freshly added Rnase inhibitors (80 U/ml)] were used to homogenize the tissues with Dounce tissue grinder (10 strokes with loose pestle and 10 strokes with tight pestle). Homogenates were transferred into an ultracentrifuge tube on top of 10 ml of fresh nuclei homogenization buffer and centrifuged at 24,000 rpm for 60 min at 4°C on a high-speed centrifuge. The nuclei pellet was resuspended in 1 ml of nuclei resuspension buffer [15 mM Hepes pH 7.4, 15 mM NaCl, 60 mM KCl, 2 mM MgCl_2_, 3 mM CaCl_2_, and freshly added Rnase inhibitors (80 U/ml)] and counted using a hemocytometer. The nuclei were centrifuged at 800 g for 10 min at 4°C and resuspended at a concentration of 500 to 1000 nuclei/μl for 10x Genomics Chromium loading. The single nucleus (sn)RNA-seq libraries were prepared by the Chromium Single Cell 3′ Reagent Kit v3.1 chemistry according to the manufacturer’s instructions (10x Genomics) and sequenced using Illumina NovaSeq 6000 S4 at the Yale Center for Genome Analysis (YCGA).

### Single nucleus transcriptome analysis – Data alignment

For single nucleus snRNA-seq of mouse brain nuclei, we used CellRanger (version 7.1.0; 10x Genomics) to align the reads against the mouse mm10 reference genome provided by CellRanger with “include-introns” as an option. The filtered count matrices generated by the software were used for further analysis.

### Single nucleus transcriptome analysis - Parameter selection for clustering analysis

Gene count matrices were analyzed by R package Seurat (version 4) ^40^ following standard protocol. Briefly, cells with low feature count (< 200) and high mitochondrial gene expression (>10%) were removed. The data was then normalized using *NormalizeData* function and highly variable genes were identified using the *FindVariableFeatures* function with the “vst” selection method and the number of features set to 2,000. Data from different genotypes was integrated using *FindIntegrationAnchors* and *IntegrateData* functions, and scaled using the *ScaleData* function with default parameters. The scaled data matrix was used for dimensionality reduction and clustering. RunPCA function was used to perform principal component analysis (PCA, *RunPCA* function in Seurat) computing the top 50 principal components. Significant principal components were identified using elbowplot and 20 principal components kept for the integrated dataset).

Clustering of the gene expression data from individual nuclei into transcriptionally similar clusters was done using a *k*-nearest neighbor graph using the *FindNeighbors* function. To cluster the nuclei, the *FindClusters* function was applied. The best resolution for clustering were found visually using R package *clustree.* We used Uniform Manifold Approximation and Projection (UMAP) for cluster visualization. The cluster markers were identified by running the *FindAllMarkers* function with default parameters. One cluster was identified and filtered out as a doublet based on high UMI and gene number per nucleus and expression of markers from more than one cell type. Cell clusters were annotated in a semi-automated approach using a combination of automatic cell annotation with Azimuth ^41^ and manual analysis of expression of known marker genes of mouse brain cell types.

### Subset analysis

For sub-clustering analysis, nuclei from the OPCs and oligodendrocytes identified in primary clustering were extracted, reclustered, and annotated based on subtype marker genes from Marques *et al.,* 2016. Small contaminating clusters expressing markers of other cell types were removed. One subset was built for the P30 control, P30 *Ng2-cre*^+^ *Prkci* ^f/f^ and P30 *Ng2-cre*^+^ *Prkcz* ^f/f^, P60 control, and P96 control to decipher the development, samples were downsampled to equalize cell numbers. Another subset was created for the *Plp1-cre*^ERT+^ samples and the cuprizone experiments.

### Identification of DEGs

DEG identification was performed using the *FindMarkers* function from the Seurat package, applying the non-parametric Wilcoxon rank-sum test with default settings. Significant DEGs were filtered for downstream pathway analysis based on criteria of log2(Fold Change) > 0.25 and adjusted p value < 0.05.

### Gene set score analysis

Gene set scores were derived via the *AddModuleScore* function in the Seurat package. The myelination score was computed using genes from the “Myelination” pathway, as defined in Metascape^42^. A signature for demyelination-associated oligodendrocytes (DOL) was developed based on marker genes from Hou *et al.*, 2023. Visualization of these computed scores was accomplished using the *FeaturePlot* function in Seurat. Scores for cell cycle were generated based on expression of genes associated with S-phase or G2/M-phase using Seurat *CellCycleScoring* function.

### Pathway enrichment analysis

Pathway enrichment analysis was conducted using Metascape ^42^, identifying overrepresented pathways in gene lists derived from Gene Ontology (GO) Biological Processes, WikiPathways, Reactome Gene Sets, and KEGG Pathway databases. The list of DEGs served as the input and the default settings of Metascape were employed for the analysis.

### Key resources table

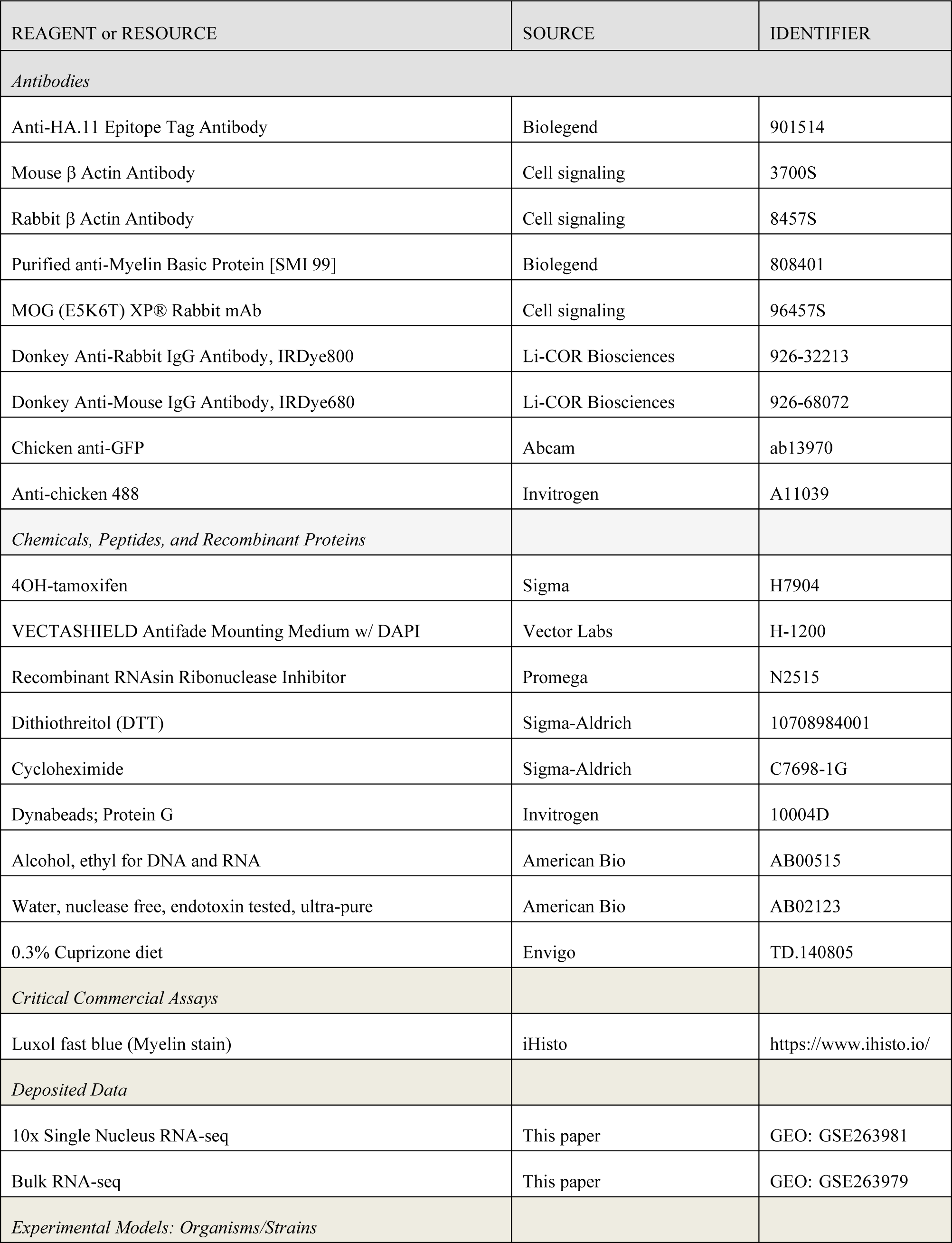
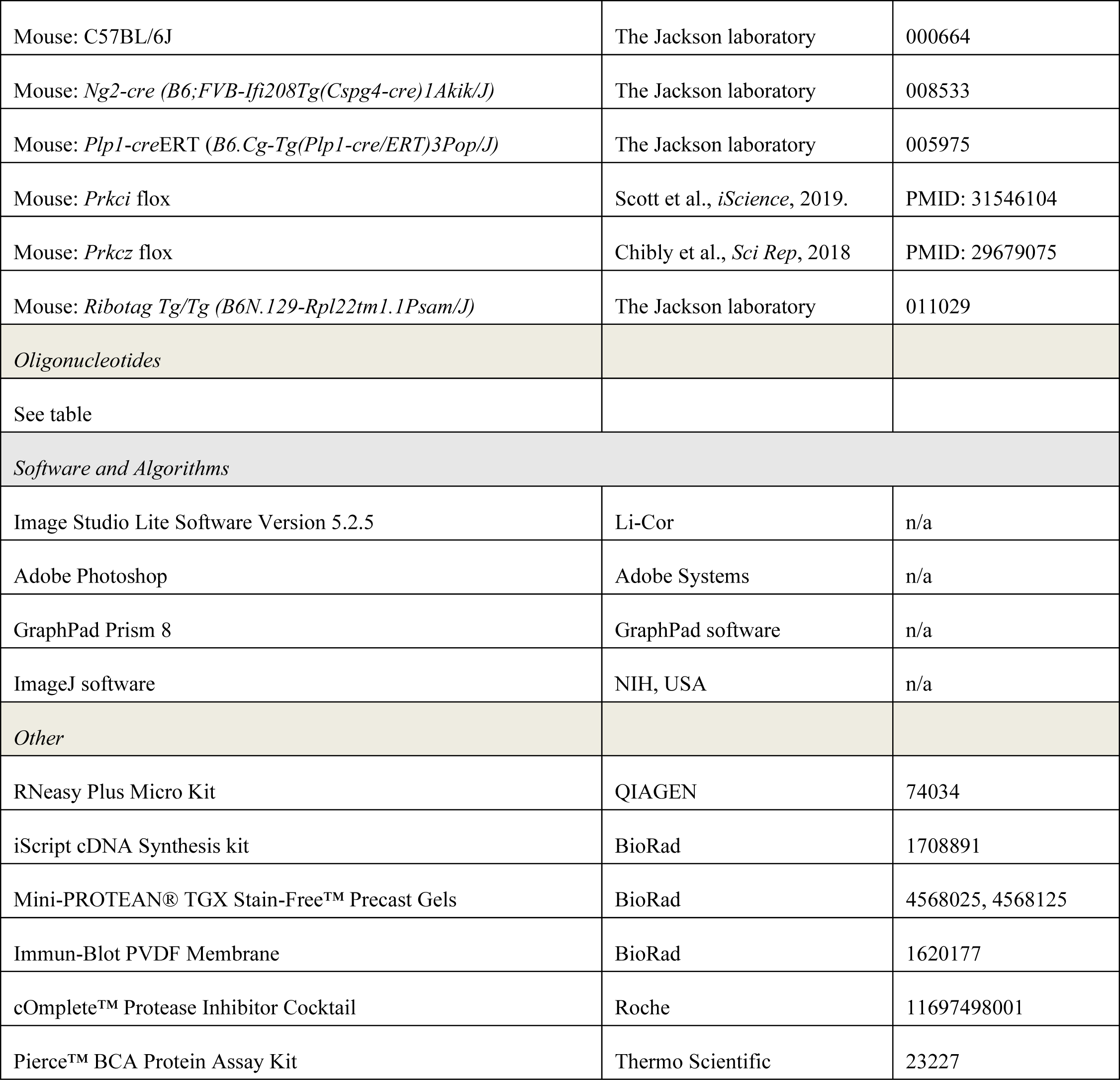

## Figures and Figure Legends

**Extended Figure S1:**
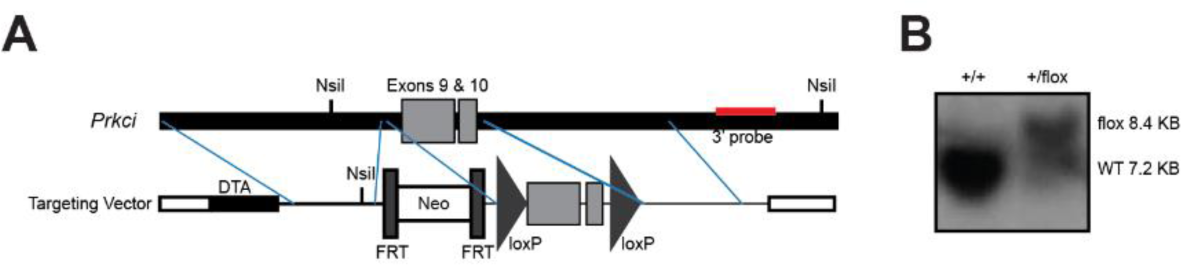
Generation of *Prkci* _f/f_ mice. **A.** Cloning strategy for the generation of *Prkci* ^f/f^ mice targeting the exons 9 and 10. **B.** Identification of the *Prkci* floxed allele by southern blot making use of a 3’ probe indicated in **A**.

**Extended Figure S2:**
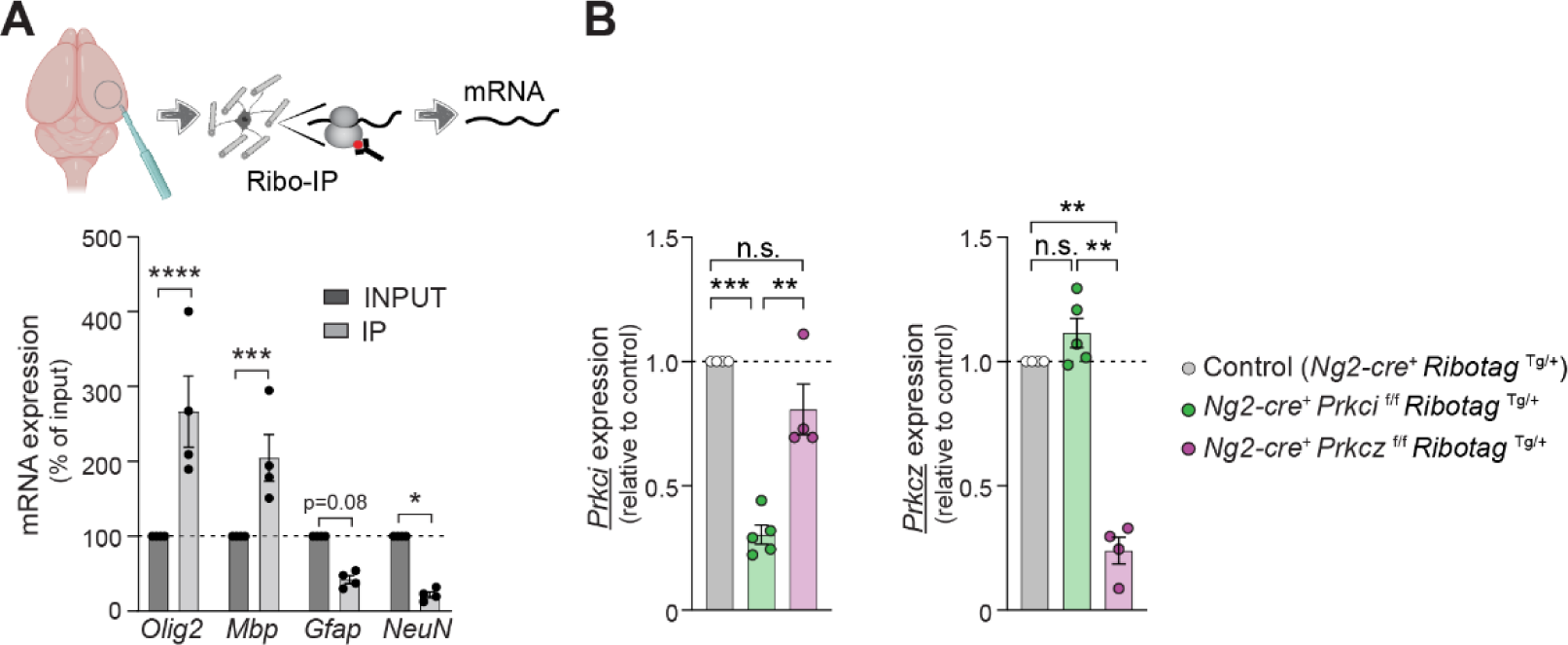
Genetic ablation of *Prkci* and *Prkcz* in NG2 lineage cells. **A.** Cerebral cortices obtained from *Ng2-cre*^+^ Ribotag ^Tg/+^, *Ng2-cre*^+^ *Prkci* ^f/f^ Ribotag ^Tg/+^ and *Ng2-cre*^+^ *Prkcz* ^f/f^ Ribotag ^Tg/+^ at P30 were processed for Ribotag immunoprecipitation. Immunoprecipitation specificity was confirmed by analysis of the expression of cell type-specific mRNA by qPCR in INPUT and IP samples. Data is shown as mean ± SEM, n=4 mice. *p<0.05, ***p<0.001, ****p<0.0001 *vs.* INPUT by 2-way ANOVA. **B.** qPCR for mRNA expression of *Prkci* and *Prkcz* in cerebral cortex at P30. Data is mean ± SEM, n=4 mice/ genotype. Non-significant (n.s.), **p<0.01, ***p<0.001 by ANOVA, followed by Tukey’s test.

**Extended Figure S3:**
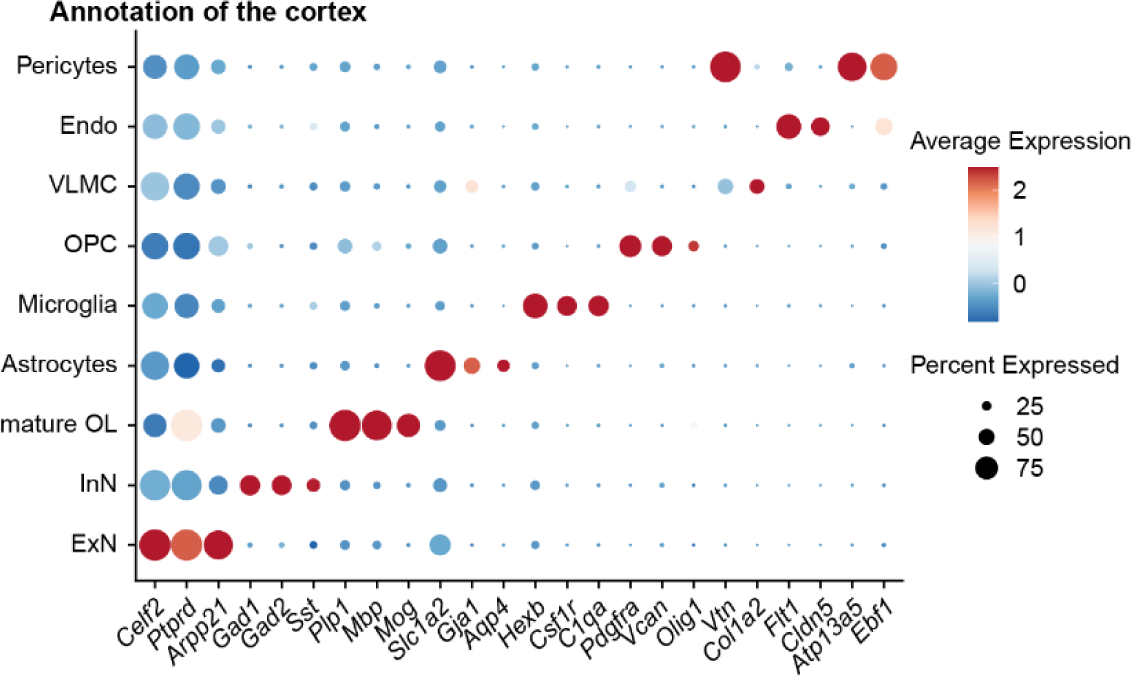
Cell-type specific gene expression for populations shown in Figure 3. Dot plot showing expression of cell-type specific genes in all nuclei clusters. ExN: excitatory neurons, InN: inhibitory neurons, mature OL: mature oligodendrocytes, OPC: oligodendrocytes precursor cells, VLMC: vascular and leptomeningeal cells, Endo: endothelial cells.

**Extended Figure S4:**
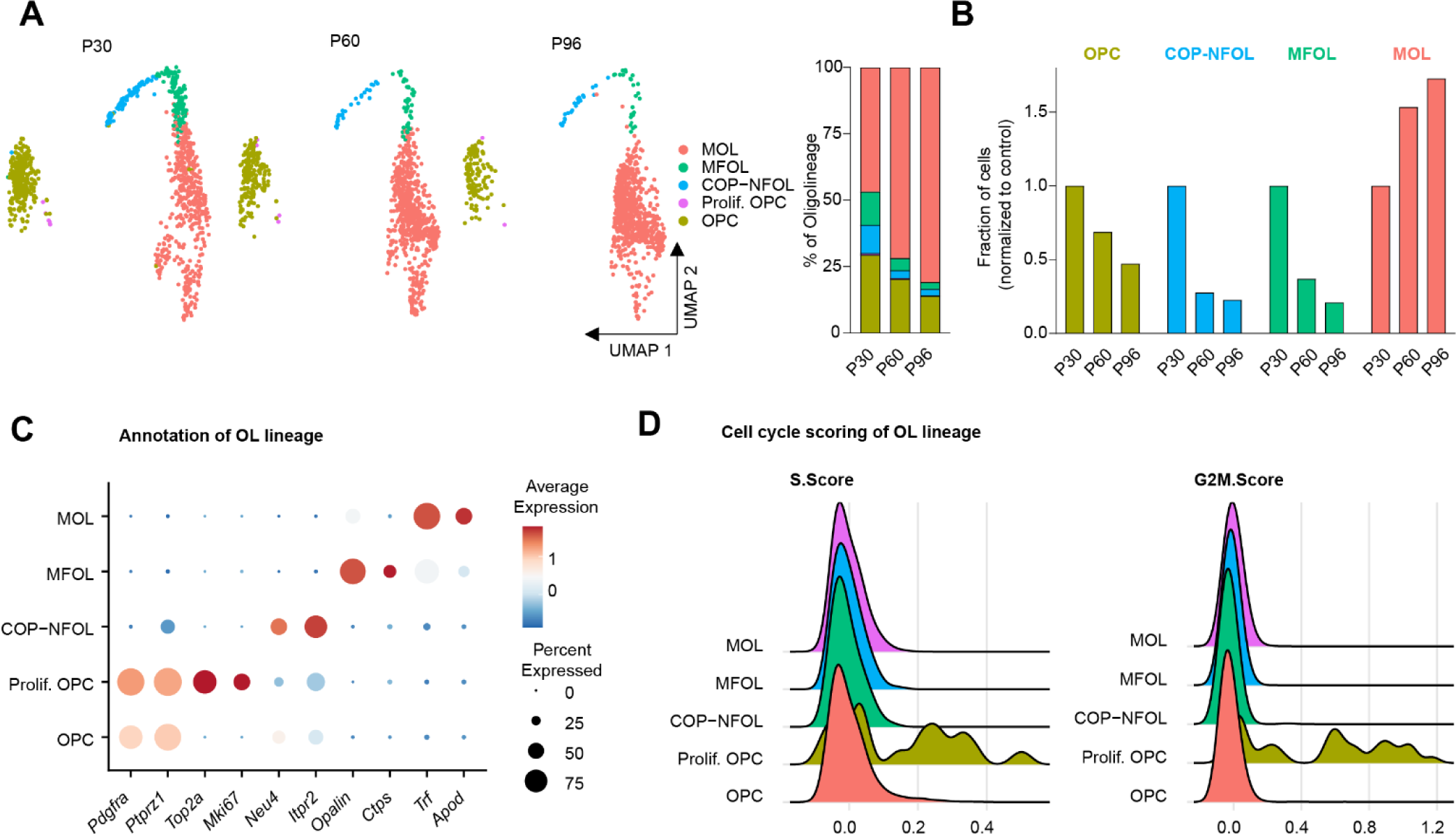
Population changes in oligo lineage from postnatal day 30 to postnatal day 96. **A.** UMAP plot of gene expression in single nuclei classified as different stages of oligodendrocyte lineage from control mice at P30, P60 and P96. Percentage nuclei representing different stages of oligodendrocyte lineage for each mouse age group are plotted on the right. **B.** Relative abundance of nuclei at different stages of oligodendrocyte lineage in P60 and P96 mice normalized to P30 mice. **C.** Dot plot showing expression of marker genes of oligodendrocyte precursor cells (OPC; *Pdgfra, Ptprz1*), proliferating OPC (Prolif. OPC: *Top2a, Mki67*), committed oligodendrocyte precursors and newly formed oligodendrocytes (COP-NFOL; *Neu4, Itpr2*) myelin forming oligodendrocytes (MFOL; *Opalin, Ctps*), mature oligodendrocytes (MOL; *Trf, Apod*) in different oligodendrocyte-lineage subclusters. **D.** Scores for cell cycle were generated based on expression of genes associated with S-phase or G2/M-phase. Ridge plot showing expression of these sets of genes in oligodendrocyte-lineage subclusters.

**Extended Figure S5:**
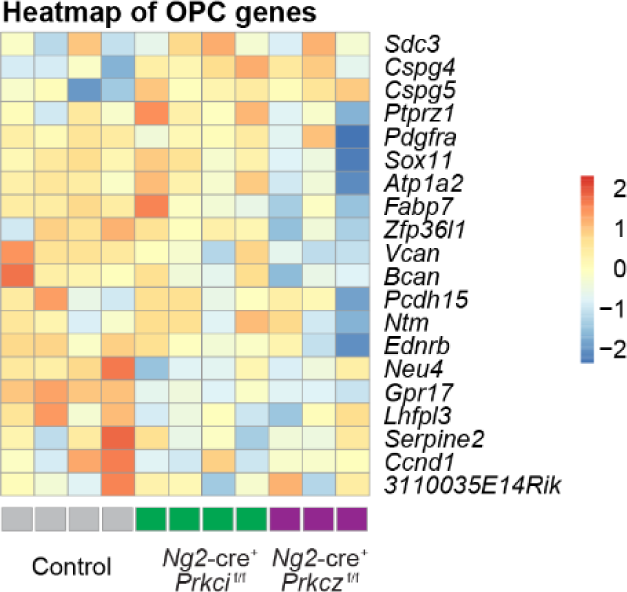
Expression of OPC genes in the cerebral cortex of control and *Ng2*-specific *Prkci* or *Prkcz* mice. Heatmap depicting expression of genes expressed by OPC (based on Marques S*. et al.* 2016). Z-score transformed gene expression values are shown, color-coded according to the color-bar on the right.

**Extended Figure S6:**
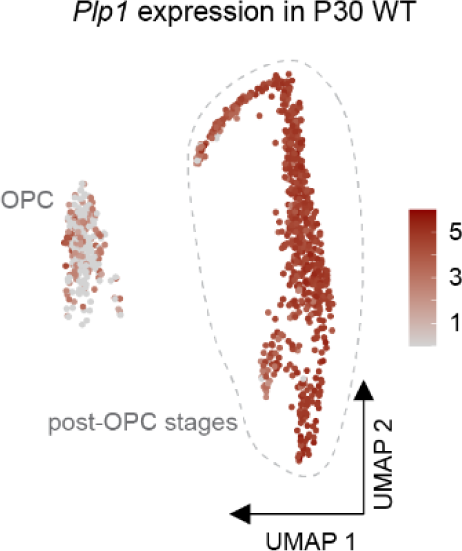
Feature plot for *Plp1* expression in indicated oligo lineage populations in WT, P30.

**Extended Figure S7:**
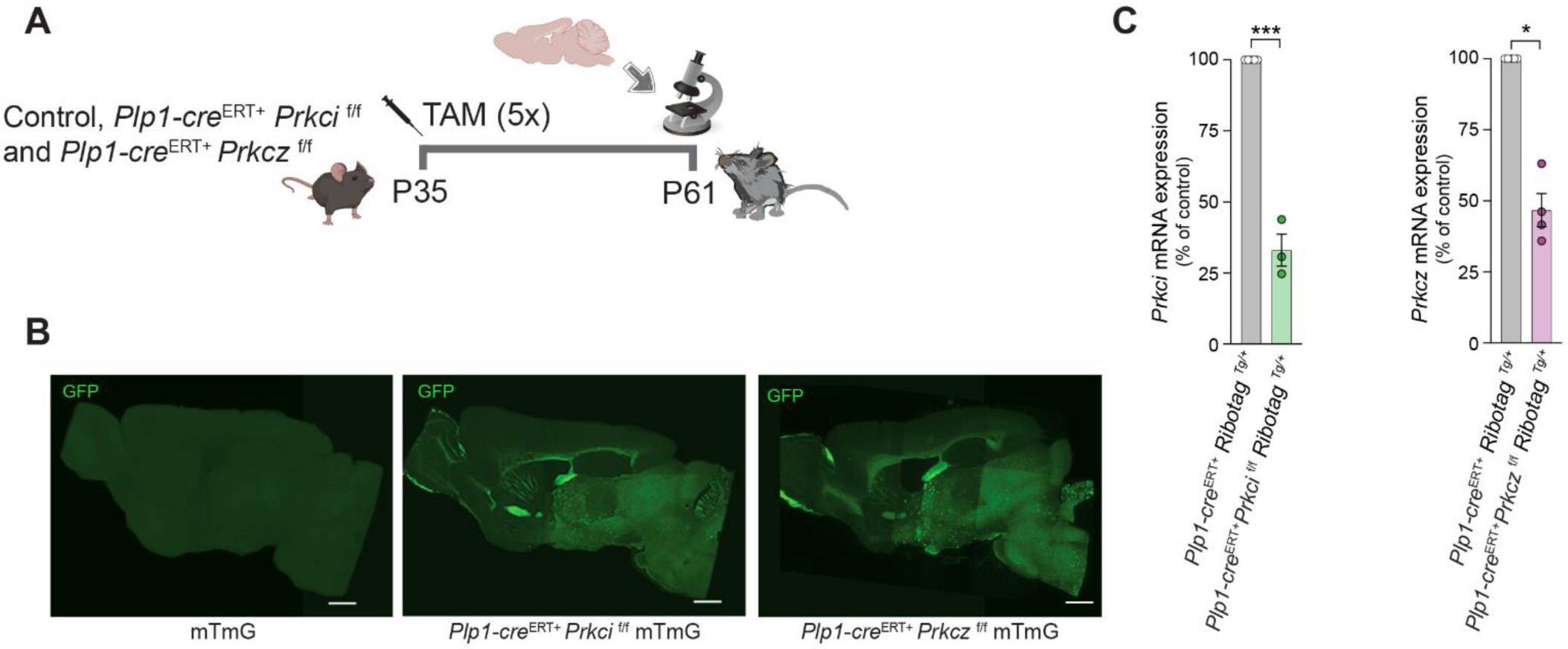
*Prkci* is dispensable for later stages of myelination of the optic nerve. **A.** Schematic depicting administration of tamoxifen (5 consecutive days, 1 mg/day, i.p.) to mTmG, *Plp-cre*^ERT+^ *Prkci* ^f/f^ mTmG, *Plp-cre*^ERT+^ *Prkcz* ^f/f^ mTmG at P35 for 5 days, sample collection was performed at 3 weeks after tamoxifen administration. **B.** *Plp-cre*^ERT+^ *Prkci* ^f/f^ and *Plp-cre*^ERT+^ *Prkcz* ^f/f^ mice were crossed to mTmG reporter mice. Tamoxifen was administered to induce activation of cre-recombinase (5 consecutive days, 1mg/day, i.p.). Cre-dependent GFP expression was analyzed by fluorescence microscopy 3 weeks after tamoxifen delivery. Representative images of tamoxifen-induced GFP expression are shown. **C.** *Plp-cre*^ERT+^ Ribotag^Tg/+^, *Plp-cre*^ERT+^ *Prkci* ^f/f^ Ribotag^Tg/+^ and *Plp-cre*^ERT+^ *Prkcz* ^f/f^ Ribotag^Tg/+^mice were administered tamoxifen to induce activation of cre-recombinase (5 consecutive days, 1 mg/day, i.p.). Oligodendrocyte-specific mRNA was extracted from cerebral cortices by immunoprecipitation of tagged ribosomes 3 weeks after tamoxifen injection, and qPCR for *Prkci* or *Prkcz* transcript 2 (encoding for PKMζ) was performed. Data is shown as mean± SEM of n=3-5 mice/genotype. *p<0.05, ***p<0.001 by Student’s t test.

**Extended Figure S8:**
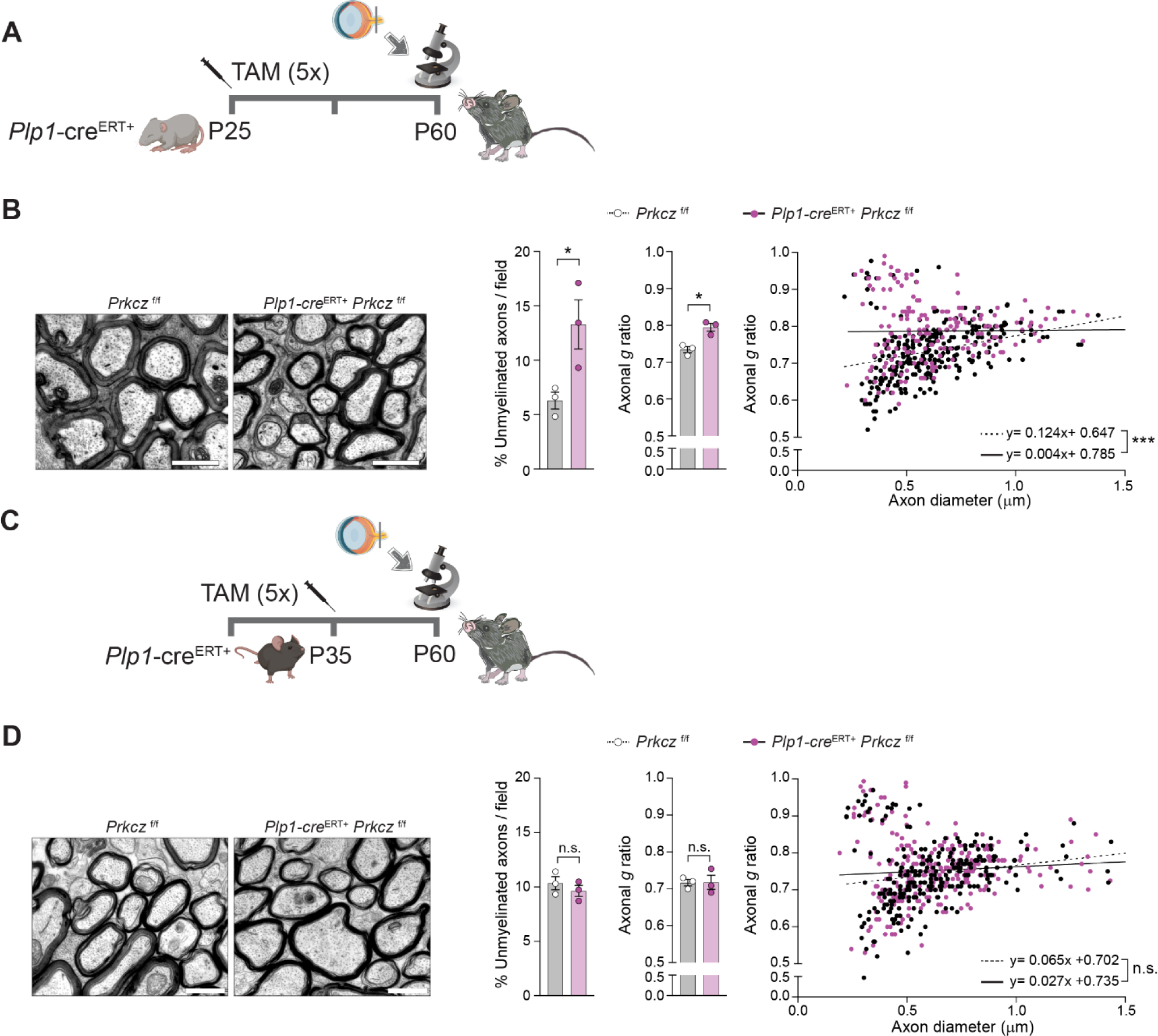
**A** and **B**. Tamoxifen was administered to *Prkcz* ^f/f^ or *Plp-cre*^ERT+^ *Prkcz* ^f/f^ mice (5 consecutive days, 1 mg/day, i.p.) starting at P25. Mice were sacrificed at P60, and optic nerves were analyzed by TEM. Representative TEM images are shown next to quantification of % unmyelinated axons in 10 fields/mouse, axonal *g* ratio in 10 fields/mouse and axonal *g* ratio *vs*. axon diameter. Data is mean ± SEM, n=3 mice/ genotype. *p<0.05 by Student’s t test. Linear regression slopes are significantly different (p= 0.0002). Scale bars = 1μm. **C** and **D.** Tamoxifen was administered to *Prkcz* ^f/f^ or *Plp-cre*^ERT+^ *Prkcz* ^f/f^ mice (5 consecutive days, 1 mg/day, i.p.) starting at P35. Mice were sacrificed at P60, and optic nerves were analyzed by TEM. Representative TEM images are shown next to quantification of % unmyelinated axons in 10 fields/mouse, axonal *g* ratio in 10 fields/mouse and axonal *g* ratio *vs*. axon diameter. Axons were determined to be unmyelinated when surrounded by a single layer of membrane/ myelin sheath. Data is mean ± SEM, n=3 mice/ genotype. Non-significant (n.s.) by Student’s t test. Linear regression slopes are not significantly different (p= 0.2157). Scale bars = 1μm.

**Table S1.**
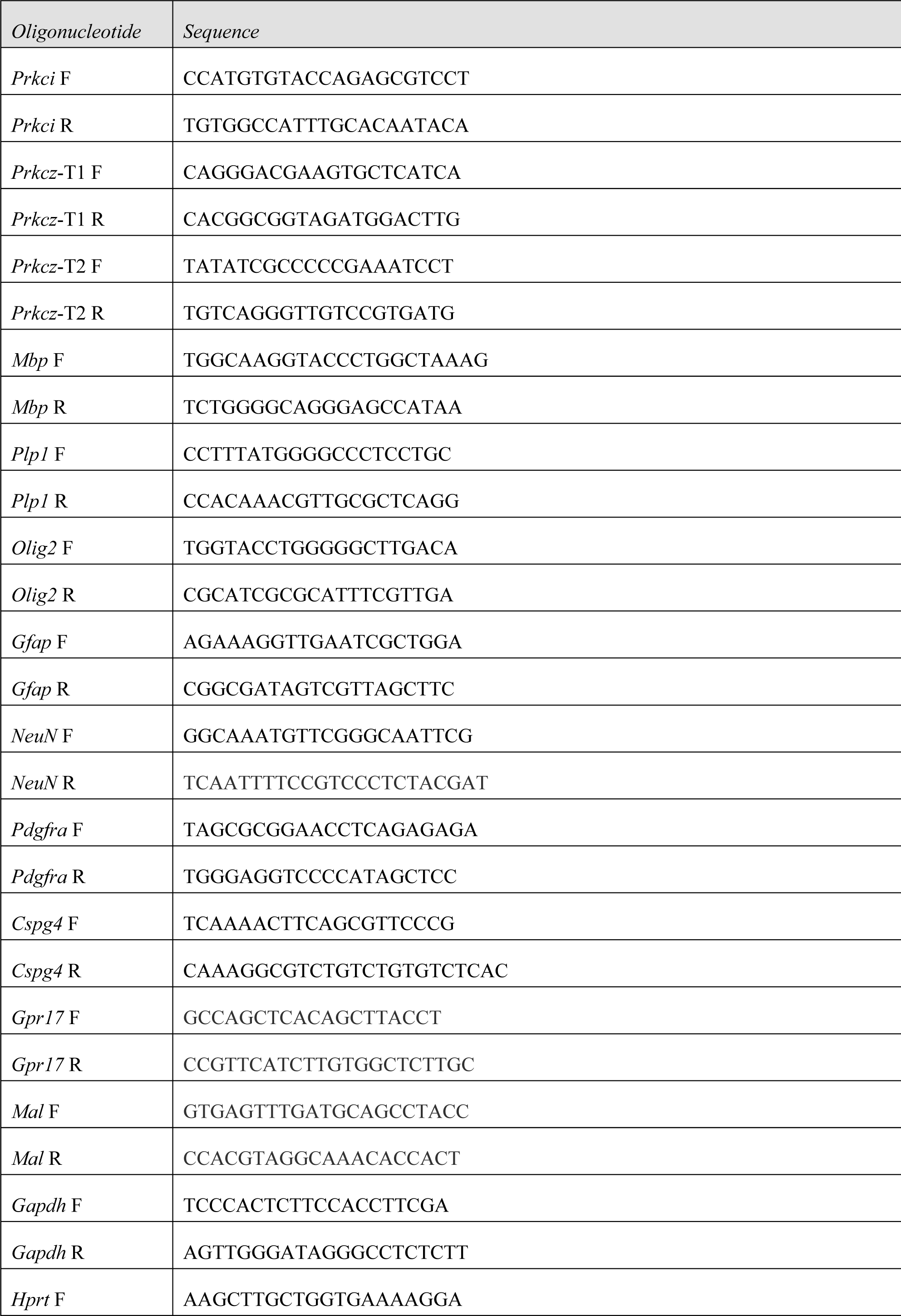
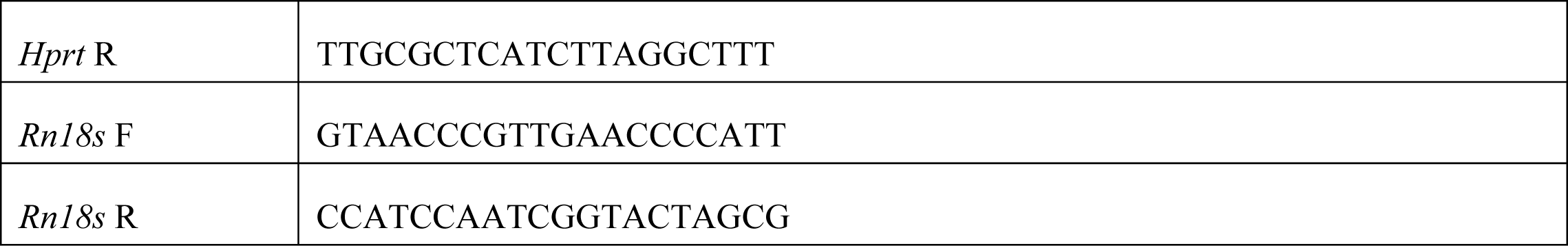
list of oligonucleotides and sequences used in this study.

